# PHA-4/FoxA controls the function of pharyngeal and extrapharyngeal enteric neurons of *C. elegans*

**DOI:** 10.1101/2025.08.13.670029

**Authors:** Zion Walker, Wen Xi Cao, Eduardo Leyva-Diaz, Mayeesa Rahman, Surojit Sural, Michelle A. Attner, Oliver Hobert

## Abstract

FoxA transcription factors pattern gut tissue across animal phylogeny. Beyond their early patterning function, little is known about whether they control the terminal differentiation and/or function of the fully mature enteric nervous system, the intrinsic nervous system of the gut. We show here that the expression and function of the sole *C. elegans* FoxA homolog, PHA-4, reaches beyond its previously described pioneer factor roles in patterning the foregut. Through the engineering of neuron-specific *cis*-regulatory alleles, Cre-mediated cell-specific knockouts and degron-mediated, temporally controlled PHA-4/FoxA removal in postmitotic neurons, we found that PHA-4/FoxA is required not only to initiate the terminal differentiation program of foregut-associated enteric neurons, but also to maintain their functional properties throughout the life of the animal. Moreover, we discovered novel sites of expression of PHA-4/FoxA in extrapharyngeal enteric neurons that innervate the hindgut (AVL and DVB), a GABAergic interneuron that controls foregut function during sleep (RIS), and a peptidergic neuron, PVT, which we implicate here in controlling defecation behavior. We show that while PHA-4/FoxA is not required for the developmental specification of AVL, DVB, RIS, and PVT, it is required to enable these neurons to control enteric functions. Taken together, *pha-4* is the only transcription factor known to date that is expressed in and required for the proper function of all distinct types of enteric neurons in a nervous system.

## INTRODUCTION

The gut of adult animals, subdivided into foregut, midgut, and hindgut, is composed of a multitude of distinct cell types. These include enteric neurons that innervate gut tissue to control various aspects of gut function, most notably, the contraction of gut musculature. The enteric nervous system was initially dubbed the “second brain” of an animal to illustrate its physical separation from the CNS and its autonomous function (Gershon 1998). Similarities in the organization and function of the vertebrate enteric nervous system to the nervous system of comparatively more primitive animals such as cnidarians, have resulted in the enteric nervous system also being dubbed the “first brain”, i.e., an evolutionary antecedent of a more complex nervous system (Furness and Stebbing 2018). These evolutionary considerations, as well as the involvement of enteric nervous systems in many aspects of human health and disease (Rao and Gershon 2018), have spurred substantial interest in a better understanding of the developmental programs that specify enteric nervous systems. Notably, while early patterning mechanisms that result in the eventual specification of enteric neurons are relatively well understood in a number of distinct invertebrate and vertebrate animal systems (Hartenstein 1997; Laranjeira and Pachnis 2009; Mango 2009; Sasselli et al. 2012; Nagy and Goldstein 2017), the molecular mechanisms that operate in postmitotic enteric neurons to specify their terminal differentiation program and to maintain their functional features during the life of the animal remain incompletely understood.

FoxA-type forkhead domain transcription factors control the specification of gut tissue across animal phylogeny, ranging from protostomes to deuterostomes (Friedman and Kaestner 2006; Mango 2009; Adler et al. 2014; Annunziata et al. 2019). For example, in the nematode *C. elegans*, expression of the sole FoxA transcription factor orthologue, PHA-4, demarcates all cells in the developing embryo that generate foregut, midgut, and hindgut tissue (Azzaria et al. 1996; Horner et al. 1998; Kalb et al. 1998; Mango 2009; Niu et al. 2011; Ma et al. 2021). Previous genetic loss-of-function studies have firmly established functions of PHA-4/FoxA in all sections of the gut at different stages of embryonic development (Mango et al. 1994; Horner et al. 1998; Kalb et al. 1998; Gaudet and Mango 2002; Mango 2009; Fakhouri et al. 2010). In the fore- and hindgut, *pha-4* functions as an organ selector to control the lineage specification and morphogenesis of these tissues during embryonic patterning. The organ selector role of PHA-4 critically depends on a phylogenetically conserved pioneer factor function of PHA-4, which involves nucleosome displacement and RNA Pol II recruitment (Hsu et al. 2015).

In the midgut (intestine), *pha-4* may not have a developmental function (a notion that we probe further in this manuscript) but has a well-documented role in intestinal homeostasis (Panowski et al. 2007; Wu et al. 2018; Torzone et al. 2023). Outside the gut, PHA-4 has only been reported to be expressed in a small number of postembryonically generated cells, which share with the alimentary tract the feature of also forming tubular structures, specifically, the somatic gonad (Kalb et al. 1998; Updike and Mango 2007; Chen and Riddle 2008). PHA-4 appears to function in the proper differentiation of these cells as well (Ao et al. 2004; Updike and Mango 2007; Chen and Riddle 2008).

We set out to investigate *C. elegans* PHA-4 function because of our overall interest in studying terminal differentiation programs of enteric neurons of the foregut and the hindgut (Hobert et al. 1999; Gendrel et al. 2016; Vidal et al. 2022). Two enteric neurons, AVL (located in the ventral head ganglion) and DVB (located in the dorsorectal ganglion of the tail), which we refer to here as “<u>h</u>indgut <u>e</u>nteric <u>n</u>eurons” (HENs), innervate the hindgut of the worm to control enteric muscle contraction during defecation (McIntire et al. 1993; Avery and Thomas 1997)(**Fig.1A**). AVL and DVB are GABAergic neurons that are activated by the intestinally released neuropeptide *nlp-40*, which generates all-or-none action potentials in these neurons (Mahoney et al. 2008; Wang et al. 2013; Jiang et al. 2022). The rhythmic activation of AVL and DVB results in the GABA-dependent contraction of electrically coupled hindgut enteric muscles (stomatointestinal, anal depressor, and anal sphincter muscles), leading to the expulsion of gut contents (Thomas 1990; Jiang et al. 2022). The terminal identity features of the AVL and DVB are genetically specified by the LIM homeobox gene *lim-6* and the orphan nuclear hormone receptor *nhr-67* (Hobert et al. 1999; Gendrel et al. 2016).

**Figure 1:**
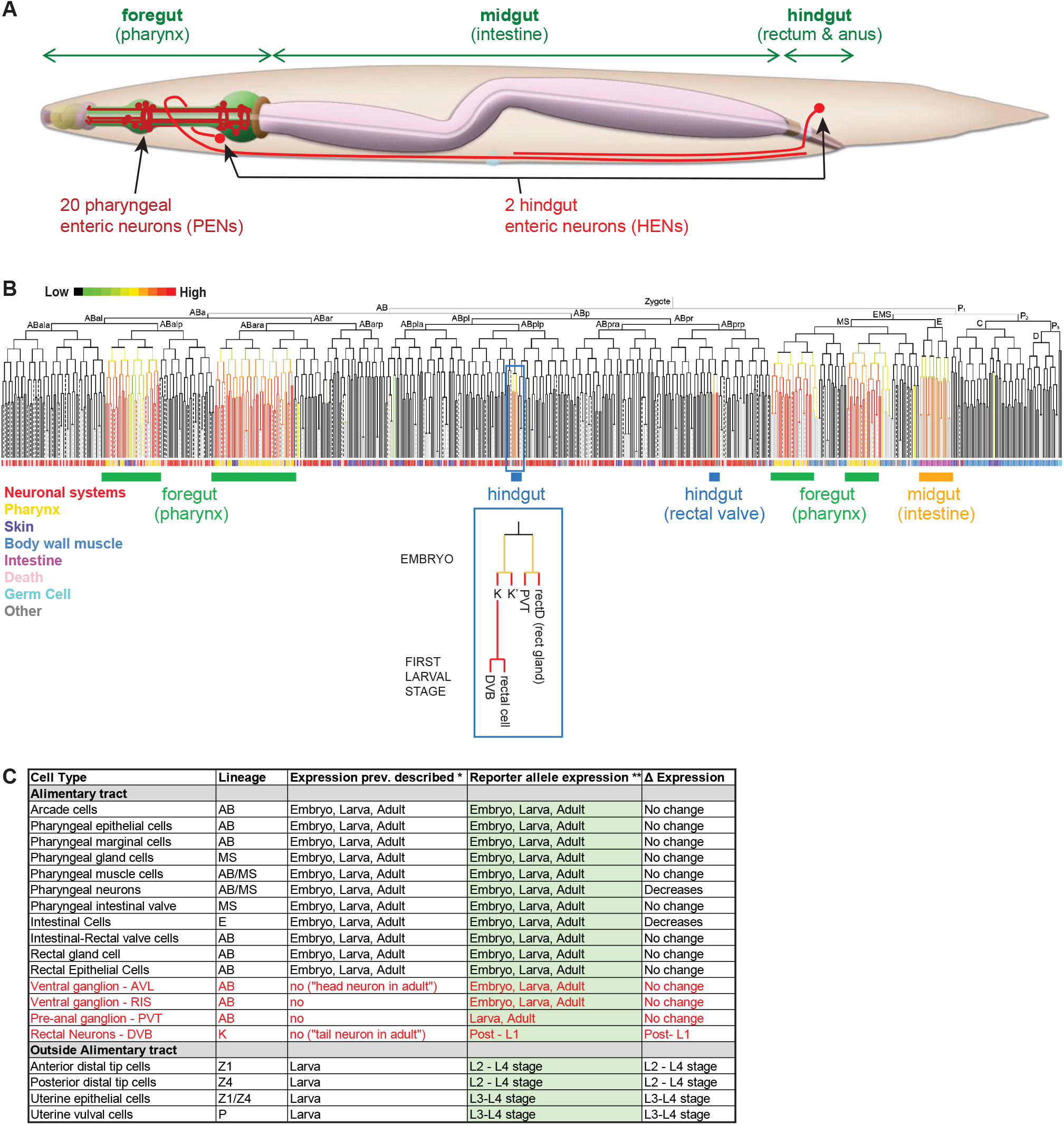
The *C. elegans* alimentary system, enteric neurons and PHA-4 expression. **A:** Schematic overview of enteric system, including schematics of the pharyngeal and hindgut enteric neurons. Adapted from *WormAtlas* (Hall and Altun 2007). **B:** Embryonic expression of PHA-4 as determined by 4D microscopy of a *pha-4* fosmid-based reporter, adapted from Ma et al., 2021. The lineage that generates PVT is highlighted to show its relation to hindgut cells (gland, hindgut-innervating DVB, rectal epithelial cells). **C:** Summary of previously reported and new expression pattern of PHA-4. * identified in previous studies using antibody staining and/or reporter transgenes (Horner et al. 1998; Kalb et al. 1998; McKay et al. 2003; Grosshans et al. 2005; Panowski et al. 2007; Chen and Riddle 2008; Riddle et al. 2016); ** this paper. “D Expression” refers to expression changes observed through stages of animal development. Newly defined sites of expression are in red.

In addition, 20 enteric neurons are located within the foregut of the worm, which we term here “<u>p</u>haryngeal <u>e</u>nteric <u>n</u>eurons” (PENs)(**Fig.1A**). These 20 neurons can be classified into 14 distinct types based on position, neurite morphology, synaptic connectivity and molecular profile (Albertson and Thomson 1976; Cook et al. 2020; Taylor et al. 2021). Like enteric neurons in all other animal nervous systems, PENs form a densely interconnected, autonomously acting network that controls foregut contraction during feeding behavior known as pharyngeal pumping (Albertson and Thomson 1976; Cook et al. 2020).

All 20 PENs require the Six-type homeodomain transcription factor CEH-34 to acquire their unique terminal properties, with CEH-34 interacting with distinct homeodomain cofactors to specify the distinct identity of individual enteric neurons (Vidal et al. 2022). We had previously reported that of the many distinct functions of PHA-4 during organogenesis and specification of distinct foregut tissue, one is the proper induction of CEH-34 expression in foregut enteric neurons (Vidal et al. 2022). As we describe here in more detail, PHA-4 is not only expressed during embryonic induction of CEH-34 expression, but its expression is maintained in all enteric neurons throughout postdevelopmental stages into adulthood. Surprisingly, we also discovered previously unnoted and continuous PHA-4 expression in the extrapharyngeal HENs (AVL and DVB), as well as in two other extrapharyngeal neurons, RIS and PVT. RIS was previously shown to control foregut-associated behavior (pharyngeal pumping)(Steuer Costa et al. 2019), and we show here that PVT controls hindgut-associated behavior (defecation).

Postmitotic, postdevelopment expression of PHA-4 in these various enteric function-controlling neuron classes indicates that PHA-4 may have late functions after its earlier functions in foregut organogenesis. Through the engineering of neuron-specific cis-regulatory alleles, Cre-mediated cell-specific knockout, and degron-mediated temporally controlled PHA-4 removal, we show here that PHA-4 function extends beyond an early pioneer function to later postdevelopmental roles within pharyngeal enteric neurons, as well as in the extrapharyngeal neurons that control enteric function. Our analysis reveals that PHA-4 represents the only currently known transcription factor whose expression and function covers the entire set of enteric neurons in an animal nervous system.

## RESULTS

### Expression pattern of *gfp*-tagged *pha-4*

The expression pattern of *pha-4* throughout the entire animal has been previously analyzed using multicopy reporter transgenes and PHA-4 antibody staining (Horner et al. 1998; Kalb et al. 1998; McKay et al. 2003; Grosshans et al. 2005; Panowski et al. 2007; Chen and Riddle 2008; Riddle et al. 2016). 4D live imaging analysis of a PHA-4 fosmid-based reporter further refined the pattern of PHA-4 expression onset during embryogenesis (Ma et al. 2021) (**Fig.1B,C**). One notable aspect of these previous analyses was the observation of PHA-4 expression in a previously unidentified head and tail neuron in adult animals (Kalb et al. 1998; Panowski et al. 2007; Chen and Riddle 2008)(**Fig.1C**). To confirm all these previous patterns using an independent reagent and to identify the nature of the PHA-4 expressing head and tail neurons, we tagged the endogenous *pha-4* locus with *gfp* using CRISPR/Cas9 genome engineering. The locus was tagged at the 3’end to capture all three isoforms of the *pha-4* locus (**Fig.2A**). Resulting *pha-4(ot946[pha-4::gfp])* animals are viable and appear superficially wildtype, indicating that tagging does not obviously affect gene function.

**Figure 2:**
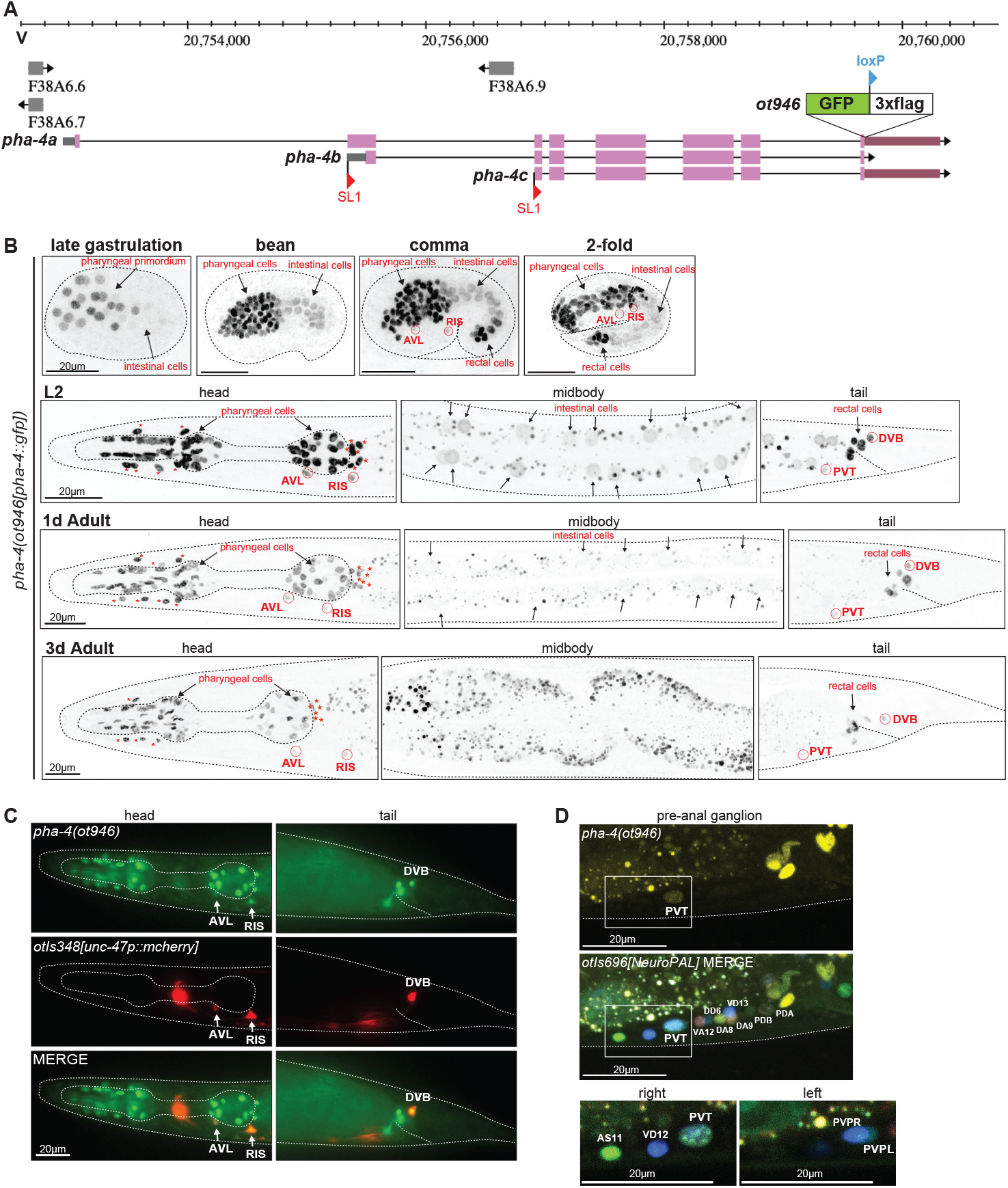
Expression pattern of a *gfp*-tagged *pha-4* locus. **A:** *pha-4* locus schematic on Chromosome V, showing the *pha-4* CRISPR/Cas-9-engineered reporter allele *pha-4(ot946[pha-4::gfp::loxP::3xflag)*, indicating the location of the GFP, loxP and 3xFLAG cassette insertion. The existence of different isoforms is inferred from the identification of different SL1-trans-spliced transcripts (Azzaria et al. 1996). **B:** Expression of *pha-4*(*ot946*) *gfp* reporter allele over the course of development, showing expression in the enteric system throughout different embryonic and postembryonic stages, including adulthood. Expression in the intestine strongly decreases in adulthood. Asterisks (*) indicate the arcade cells. Carets (^) indicate the pharyngeal-intestinal valve cells. Overlays with a panneuronal reporter are shown in Fig.S1A. **C:** *pha-4* is expressed in GABAergic neurons AVL, RIS, and DVB. Images of an adult worm show co-localization of *pha-4::gfp* expression with a *unc-47prom::mcherry* reporter transgene (*otIs348*) in two neurons situated on the ventral ganglion in the head (AVL and RIS), and one neuron on the dorsal side in the tail (DVB). **D:** *pha-4::gfp* is expressed in the pre-anal ganglion neuron PVT. Magnified images of the pre-anal ganglion of an adult worm show co-localization of *pha-4*(*ot946*) expression with PVT as identified by NeuroPAL (*otIs696)*. NeuroPAL cell IDs of neighboring neurons are also annotated.

Expression of this *pha-4* reporter allele precisely recapitulated the previous reported sites of expression of PHA-4 throughout the entire alimentary system, from the arcade cells which attach the foregut to the mouth, to all cells of the foregut, midgut (intestine) and cells associated with the hindgut (**Fig.2B**). Expression initiates at the end of gastrulation and is evident throughout the pharyngeal primordium and the midgut (**Fig.2B)**. While expression in the midgut becomes significantly weaker over the course of embryonic and postembryonic development, *pha-4* expression in cells of the fore- and hindgut remain strong into adulthood (**Fig.2B; Fig.S1A**), a possible consequence of transcriptional autoregulation of the *pha-4* locus (Kaltenbach et al. 2005).

While previous studies noted expression only in an unidentified “head neuron” and a “rectal neuron” (Kalb et al. 1998; Panowski et al. 2007; Chen and Riddle 2008), we noted a total of four extra-pharyngeal neurons that express *pha-4(ot946)* in the adult. Based on position, and with the use of molecular landmarks, we identified these as the GABAergic RIS, AVL and DVB neurons and the peptidergic PVT tail neuron (**Fig.2B-D**). Onset of PHA-4::GFP expression in RIS, AVL and rectal cells, including PVT is observed in the comma stage embryos shortly after these neurons have been generated, while expression in DVB becomes apparent at the late L1 stage, when DVB is born (**Fig.2B**). All four neurons are known to require a terminal selector transcription factor, the LIM homeobox gene *lim-6*, for their proper differentiation (Hobert et al. 1999; Aurelio et al. 2003; Gendrel et al. 2016); we found that in *lim-6* null mutant adult animals, *pha-4* reporter allele expression is still present in these neurons, but notably weaker, indicating that *lim-6* is required to sustain proper *pha-4* expression expression (**Fig.S1B**).

### *pha-4-*expressing extrapharyngeal neurons control enteric behavior

Three of the four PHA-4-expressing, extrapharyngeal neurons have been previously linked to the control of gut function. The head neuron RIS, whose single axon extends around the isthmus of the pharynx (**Fig.3A**), controls foregut function, as inferred from optogenetic activation of RIS affecting serotonin-induced pharyngeal pumping rate (Steuer Costa et al. 2019). We sought to corroborate the involvement of RIS in controlling pharyngeal pumping by silencing RIS using a transgene of *flp-11*-promoter driving expression of a histamine-gated chloride channel (HisCl) (Grubbs et al. 2020). We found that histamine-induced silencing of RIS in freely moving animals decreases pharyngal pumping behavior, while not having any impact on hindgut-controlled defecation behavior (**Fig.3D**).

**Figure 3:**
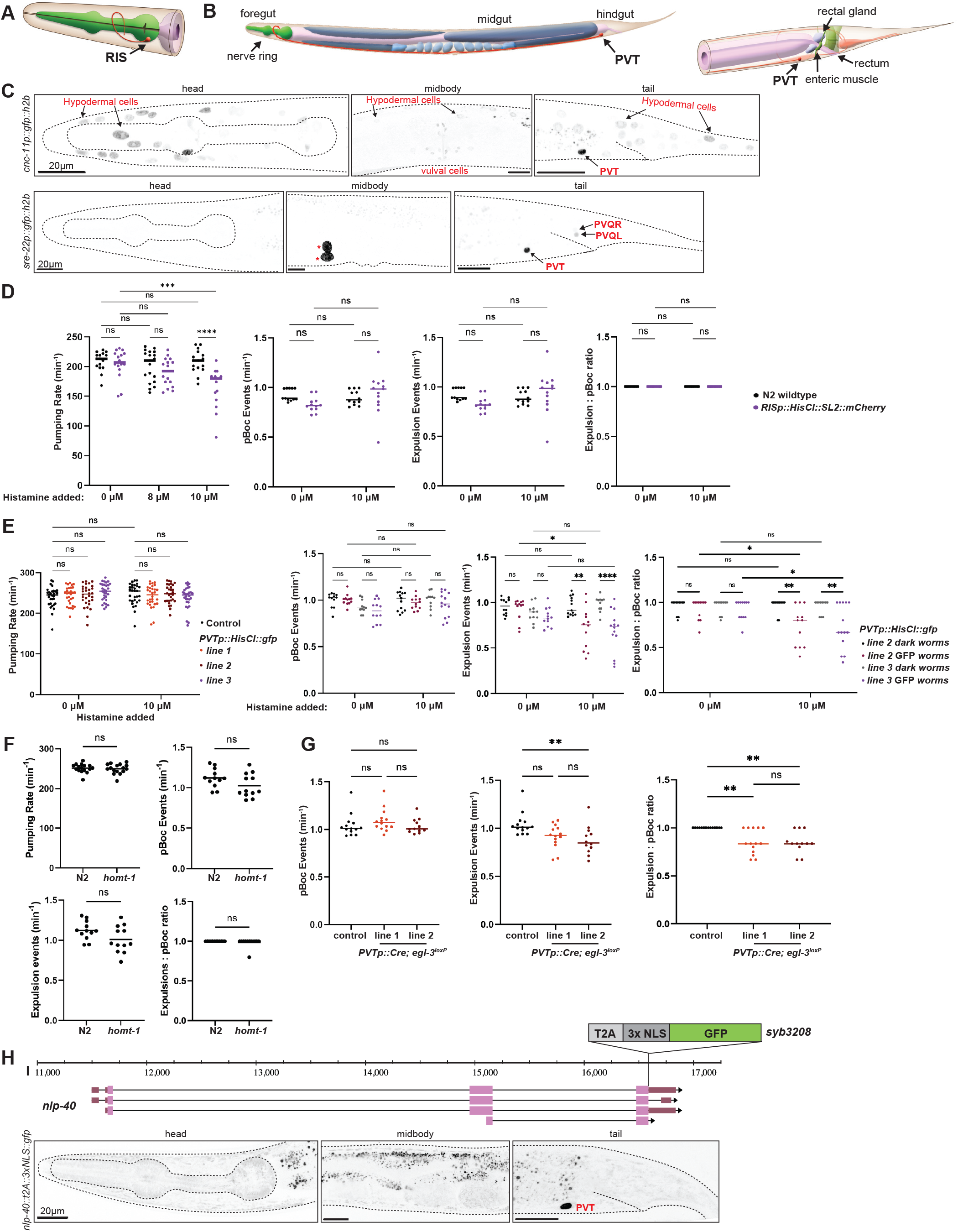
Effect of RIS and PVT inhibition on enteric behaviors. **A, B:** Schematic showing the anatomical location and morphology of the unilateral RIS neuron (A) and the PVT neuron (B), taken from *WormAtlas* at www.wormatlas.org (Hall and Altun 2007). **C:** Establishing Cre-driver lines with PVT specificity. Representative images of animals assayed in day 1 adult stage. Expression of a *cnc-11* promoter fusion (*otEx8345*) (TOP), and *sre-22* promoter fusion *(otEx8333)* (BOTTOM) in the head, midbody, and tail. Expression in PVT and a few other cell types is annotated. Asterisks indicate the expression of the co-injection marker *unc-122p::gfp* in coelomocytes. Expression was confirmed in 14 array positive animals of *otEx8333* animals, and 25 array positive *otEx8345* animals. **D**. Histamine-mediated silencing of RIS impacts adult pumping, but not defecation behavior. Pharyngeal pumping in RISp::HisCl expressing worms (*qnEx643*). Pharyngeal pumping was observed in the absence of histamine (0 μM), and at 8 μM and 10 μM concentrations of histamine. Frequency of pBocs, frequency of expulsion, and the Expulsion:pBoc ratio in RISp::HisCl expressing adult worms treated with histamine were assayed and individually plotted. **E:** Histamine-mediated silencing of PVT impacts defecation, but not pharyngeal pumping behavior. Three PVTp::HisCl array-expressing lines (*otEx8336 otEx8337 otEx8338*; line 1, line 2 and line 3) were assayed. Pharyngeal behavior was observed at 0 μM and 10 μM concentrations of histamine (LEFT). Frequency of pBocs, frequency of expulsion, and expulsion:pBoc ratio in adult worms of two lines expressing PVTp::HisCl *(otEx8336* and *otEx8337*; line 2 and line 3) treated with histamine were assayed and individually plotted (RIGHT). Dark worms are non-array carrying siblings from the PVTp::HisCl lines (*i*.*e*. controls). **F:** Enteric behaviors are unaffected in *homt-1*(*zw94*) null mutant animals. **G:** Cre-mediated removal of *egl-3* from PVT impacts adult defecation. *PVTp::Cre; egl-3*^*loxP*^ lines (*otEx8339* and *otEx8340*). **H:** *nlp-40* locus schematic on Chromosome I, showing the *nlp-40* CRISPR/Cas-9-engineered reporter allele *syb3208*, indicating the location of the reporter cassette insertion. Previously reported *nlp-40* reporter transgene expression in the intestine appears too dim to be visible with this reporter allele. **D - G**: Data points show individual worms assayed. Horizontal line represents median value of biological replicates*, **, ***, **** and ns represent P < 0.05, P < 0.01, P < 0.001, P < 0.0001 and not significant, respectively, in a Mann-Whitney test (F), and Dunn’s multiple comparison test after Kruskal-Wallis test (G), and Sidak’s multiple comparisons test after two-way ANOVA (D and E).

AVL and DVB are known hindgut-innervating enteric neurons (“HENs”) which control enteric muscle contraction during defecation (McIntire et al. 1993)(**Fig.1A**). The defecation motor program occurs in three steps: contraction of posterior body muscles (pBoc), contraction of anterior body fmoreugsutcles (aBoc), and explusion (Exp) (Thomas 1990). A pacemaker in the intestinal epithelial cells initiates the pBoc step, which acts upstream of AVL and DVB (Dal Santo et al. 1999). The Exp step requires activation of AVL and DVB to constrict hindgut enteric muscles to release gut contents through the anus. The defecation cycle is highly regular both within and across individuals over time, with well-fed young adult animals never missing a pBoc, aBoc, or Exp step (Thomas 1990).

PVT is a peptidergic interneuron in the pre-anal ganglion whose single axon extends from the tail to the isthmus of the pharynx (**Fig.3B**). PVT displays an intriguing lineage relationship with some cellular constituents of the gut (**Fig.1B**), but no function in relation to the enteric system has been previously investigated. To manipulate PVT activity, we first used scRNA data (Taylor et al. 2021) to identify potential drivers with PVT-specific expression, which we validated with reporter gene constructs (**Fig.3C**). We used the promoter of *sre-22* to drive histamine-gated chloride channel (HisCl) in PVT and found that after histamine-treatment the frequency of pBoc events in such transgenic animals was unaffected, but the rate of expulsions was significantly reduced (**Fig.3E)**. Compared to untreated PVTp::HisCl worms, histamine-treated worms expressing PVTp::HisCl exhibited significantly less expulsion events, with an average of every third pBoc event not followed by an expulsion event (**Fig.3E)**. PVT silencing does not affect pharyngeal pumping (**Fig.3E**).

PVT has also recently been shown to control locomotory quiescence during sleep like-behavior via releasing melatonin (Niu et al. 2020). We found that animals lacking the melatonin-synthesizing *homt-1* gene display no effect in defecation behavior (**Fig.3F)**, indicating that PVT may act via other signaling pathways to control defecation behavior. Since PVT expresses a host of neuropeptides (Taylor et al. 2021), we tested whether removal of the neuropeptide-processing enzyme *egl-3* in PVT would affect defecation behavior. To this end, we used a floxed *egl-3* locus (Marquina-Solis et al. 2024) and drove Cre expression with the *sre-22* driver. We found that in worms with *egl-3* removed from PVT, the frequency of pBoc events were still unaffected, but not all pBoc events were followed by expulsions **(Fig.3G)**. Unlike control animals (floxed *egl-3* animals with no Cre expression), worms with *egl-3* removed from PVT displayed both missed expulsions and failed expulsions, in which a weak contraction of the intestinal muscle was observed, but the concerted contraction of the hindgut enteric muscles and expulsion of gut contents was absent (**Supp Video 1, 2**). While the timing of the weak muscle contraction occurs roughly 3 seconds after the pBoc in the defecation cycle, which is typical when AVL and DVB are active, the weak muscle response is not sufficient to complete the expulsion of gut contents from the body (**Supp Video 2)**. The aberrant expulsion phenotype observed in worms with *egl-3* removed from PVT points towards a potential role of PVT in the neuropeptidergic activation of AVL and DVB during the Exp step of defecation.

The neuropeptide *nlp-40* regulates the calcium transients of AVL and DVB, in order to orchestrate the release of GABA onto the hindgut enteric muscles. *nlp-40* is thought to be secreted from the intestine in response to intestinal calcium waves of the pBoc and aBoc (Wang et al. 2013). Intriguingly, as per scRNA data (Taylor et al. 2021), the most strongly expressed neuropeptide expressed in PVT is *nlp-40*. We validated this observation by engineering a *T2A:gfp::h2b* cassette into the *nlp-40* locus, confirming strong expression in PVT (**Fig.3H**). We also found clear expression of the NLP-40 receptor AEX-2 in PVT using an *SL2::gfp::h2b* cassette (Ripoll-Sanchez et al. 2023), suggesting that NLP-40 release from PVT may cooperate to amplify intestinally released NLP-40 to control the defecation behavior that is disrupted in *nlp-40* mutants (Wang et al. 2013).

### *Cis*-regulatory elements driving neuronal PHA-4 expression

An analysis of *pha-4* function in PENs and HENs, particularly in regard to the control of enteric behaviors, is complicated by (a) the early lethality of *pha-4* null mutants and (b) its expression in multiple distinct cells of the alimentary tract. As we exemplified in a previous study of an essential gene (Reilly et al. 2022), the dissection of the *cis*-regulatory architecture of a genetic locus provides a potential avenue to generate *cis-* regulatory mutant alleles in which expression of a gene is lost selectively in only specific subsets of cells that normally express this gene. Previous work had shown that a genomic 7kb fragment that contains around 2kb of sequences of the longest isoform of the *pha-4* locus, as well as several exons and introns, drives *lacZ* reporter expression throughout the entire alimentary tract, while a 300bp fragment upstream of the first isoform drives expression exclusively in the midgut (**Fig.4A**)(Azzaria et al. 1996; Kalb et al. 1998). We recapitulated the intestine-specific expression with a *gfp-*based transgene that captures the region upstream of the longest *pha-4* isoform (*pha-4prom1;* **Fig.4A,B**). Interestingly, 2,242 bp of sequences upstream of the second isoform (i.e. the first intron of the large isoform; *pha-4prom2;* **Fig.4A**) drove exclusive expression of a *gfp* reporter in all PENs and HENs (AVL, DVB), as well as the RIS and PVT neurons (**Fig.4C**). We confirmed this notion by crossing these transgenic animals with red fluorophore transgenes that marks either the entire nervous system or, selectively, the GABAergic identity of the HENs (**Fig.S1C,D**). The onset of expression of the *pha-4prom2::gfp* transgene in HENs and PENs is comparable to the onset of expression of the endogenously tagged *pha-4* locus (**Fig.4C, Fig.2B**).

**Figure 4:**
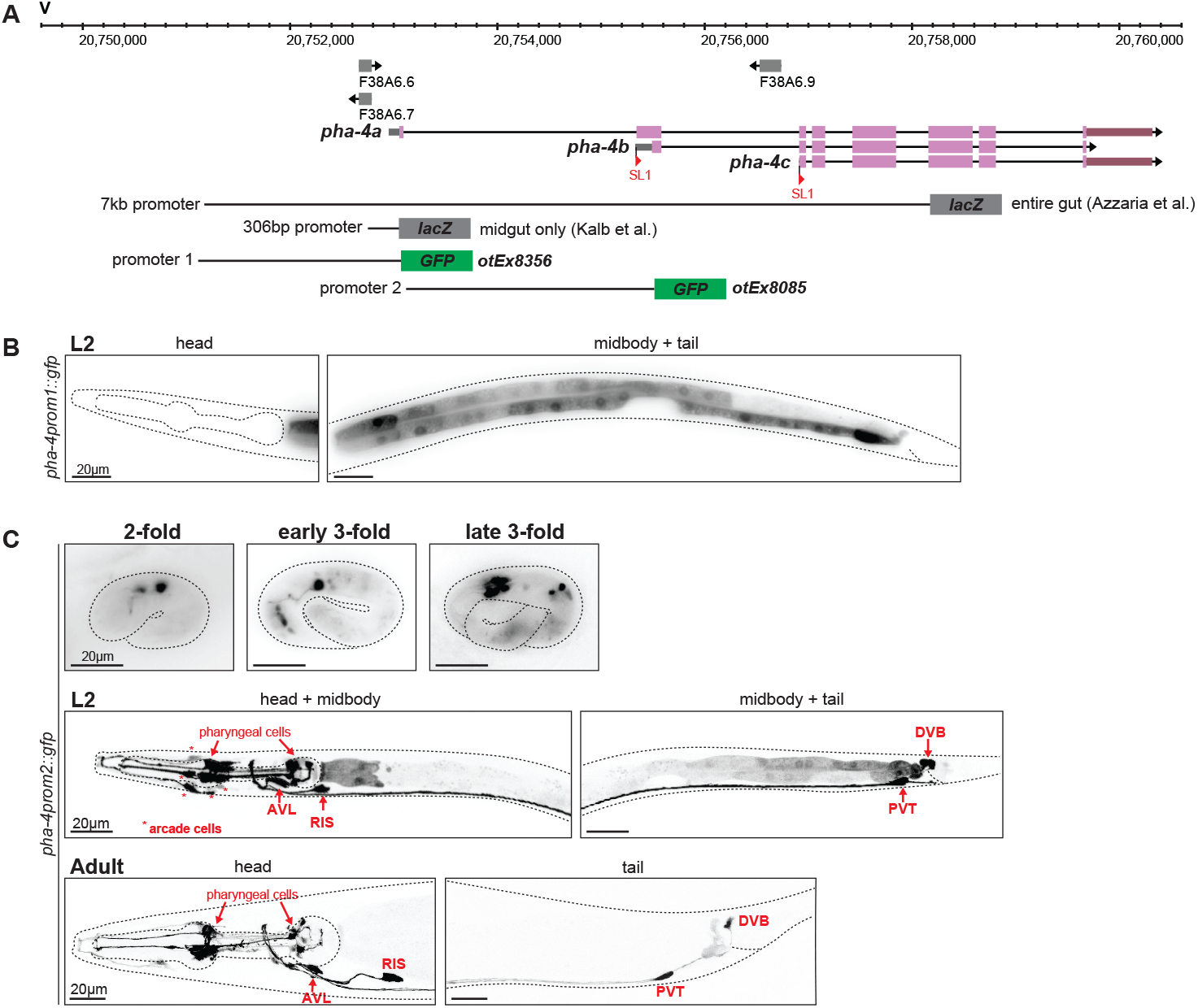
An intronic *cis*-regulatory element drives PHA-4 expression in PENs and HENs. **A:** *pha-4* locus schematic with indicated reporter gene fusions. Previously described lacZ fusions from (Azzaria et al. 1996; Kalb et al. 1998) are schematized along with two newly generated promoter fragments fused to GFP and used for expression analyses. **B:** Expression pattern of a *pha-4prom1::gfp* reporter transgene (*otEx8356*). An approximately 2kb fragment upstream of the start codon of the *pha-4a* isoform drives expression specifically in the intestinal cells. **C:** Expression pattern of a *pha-4prom2::gfp* reporter transgene (*otEx8085*). An approximately 2.4kb fragment upstream of the start codon of the second isoform, *pha-4b*, equivalent to the first intron of *pha-4a*, drives expression specifically in all enteric neurons of the pharynx, hindgut, as well as RIS and PVT. The onset of GFP expression is detected in the embryo, and expression is maintained throughout adulthood.

### Postmitotic enteric neuron expression of PHA-4 is essential for feeding and survival

Based on the *cis*-regulatory analysis described above, we used CRISPR/Cas9 to eliminate the entire first intron of *pha-4a* isoform from the endogenous *pha-4* locus that we had tagged with *gfp* (**Fig.5A**). We found that in these animals, *gfp* expression is selectively lost from PENs, HENs, RIS, and PVT (**Fig.5B**). Intriguingly, expression in these neurons appears intact initially, but is lost by the 1.5-fold stage and remains absent in L1 animals (**Fig.5B**), raising the possibility that the *pha-4prom2* region represents an autoregulatory element. To test this notion, we crossed two independent, chromosomally integrated *pha-4prom2* reporter lines into a *pha-4* null mutant background. We found that expression is mostly lost from PENs, but unaffected in two extrapharyngeal neurons, likely AVL and RIS (**Fig.5C; Fig.S1E**). We conclude that *pha-4* indeed autoregulates in many cell types, albeit apparently not in extrapharyngeal neurons.

**Figure 5:**
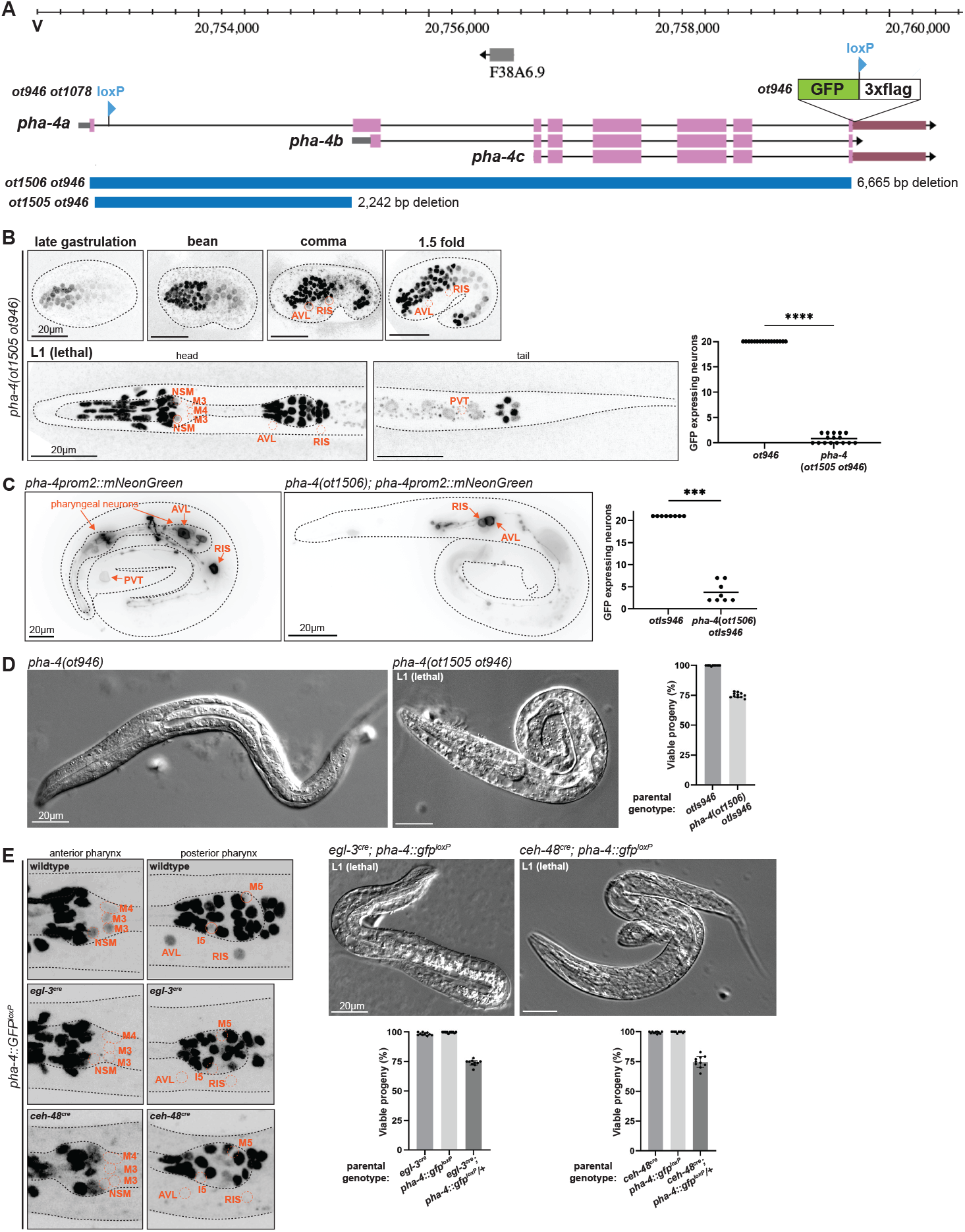
Postmitotic enteric neuron expression of PHA-4 is essential for survival. **A:** *pha-4* locus schematic on Chromosome V, showing different alleles used in this study. *pha-4(ot1078 ot946)* animals contain the C-terminal reporter cassette (*ot946;* including one loxP site), as well as an added second loxP site (*ot1078)*. The *gfp-*tagged allele was also used to either delete the first intron only, resulting in *pha-4(ot1505 ot946)* animals or to delete the entire locus, including the reporter cassette, resulting in *pha-4(ot1506)* null mutants. **B:** Expression pattern of the *pha-4 cis*-regulatory mutant allele (*ot1505 ot946*), showing lack of expression in RIS, AVL, and PVT. All other PHA-4::GFP cells in the head and tail regions of these images are non-neuronal PHA-4(+) cells. The L1 animals are shown from a dorsoventral perspective. Data points representing each individual worm assayed are plotted, and the horizontal line in the middle of the data points represents the median value of biological replicates. **** represents P < 0.0001 in a Mann-Whitney test. **C:** Expression pattern of the *pha-4* reporter transgene *pha-4prom2::mNeonGreen::PH* (*otIs946*), in wildtype or *pha-4*(*ot1506)* null mutant animals. Data points representing each individual worm assayed are plotted, and the horizontal line in the middle of data points represents the median value of biological replicates. *** represents P < 0.001 in a Mann-Whitney test.**D:** Bright field image of L1 larva expressing *gfp-* tagged *pha-4*(*ot946)* reporter allele, with wildtype morphology (left panel), and of L1 *pha-4(ot1505 ot946)* mutant lethal larva (right panel). The graph shows the survival rate of the progeny from worms heterozygous *pha-4(ot1505 ot946)* parents. One quarter of their progeny is expected to be homozygous for *pha-4(ot1505 ot946)* and we observe about one quarter of L1 arrested animals. Data points represent the percent of viable progeny of a given parent worm. 10 parent worms were assayed per condition (see Methods for more detail). **E:** Left panel: neuron-specific depletion of the floxed, *gfp-*tagged *pha-4* locus (*ot1708 ot946)* using the *Cre::egl-3*(*syb5804*) and *Cre::ceh-48*(*syb5859*) alleles was confirmed by loss of PHA-4::GFP signals in neurons of L1 lethal animals. All remaining PHA-4::GFP cells in the head region are non-neuronal PHA-4(+) cells. Right panel: Bright field images of L1 lethal larva resulting from *pha-4* removal from PENs using the *Cre::egl-3*(*syb5804*) or *Cre::ceh-48(syb5859*) alleles. Also shown is the survival rate of the progeny from worms heterozygous for *pha-4*(*ot1708 ot946*) parents in the *Cre::egl-3(syb5804*) or *Cre::ceh-48*(*syb5859*) allele backgrounds. One quarter of their progeny is expected to carry a homozygously, panneuronally deleted *pha-4* locus and we indeed observe about one quarter of L1 arrested animals. Data points represent the percent of viable progeny of a given parent worm. 10 parent worms were assayed per condition (see Methods for more detail).

Animals carrying this *cis-*regulatory allele of *pha-4* display no obvious pharyngeal morphology defects, but arrest development at the first larval stage, as scrawny, starved looking animals (**Fig.5D**). This is exactly the phenotype expected from loss of function of PENs, specifically, the M4 motor neuron, whose microsurgical removal results in exactly the same L1 arrest phenotype (Avery and Horvitz 1989).

Even though the temporal dynamics of expression of the *pha-4 cis*-regulatory allele (loss only in postmitotic neurons) and its resulting larval arrest phenotype already strongly argue for postmitotic neuronal function of *pha-4*, we sought to independently validate this notion, using an orthogonal approach. To this end, we first engineered two loxP sites into the *pha-4* locus (**Fig.5A**). We independently engineered two strains in which we inserted a *FLAG::NLS::Cre::SL2* expression cassette at the 5’end of two different genes, *egl-3* (neuropeptide-processing enzyme) and *ceh-48* (homeobox gene)**(Fig.S2;** from here on referred to as “*Cre::egl-3*” and “*Cre::ceh-48*”). We had previously shown through *gfp* tagging of these loci that both loci are expressed exclusively in all postmitotic neurons of the central, peripheral and enteric nervous systems (Leyva-Diaz and Hobert 2022). We crossed *Cre::egl-3* and *Cre::ceh-48* independently with the floxed *pha-4* locus and confirmed the depletion of *pha-4::gfp* expression in normally *pha-4-*expressing neurons (**Fig.5E**). In both resulting strains, we observed the same phenotype that we observed upon removal of the *pha-4prom2* elements: In each strain, neuronal expression of PHA-4::GFP is lost by the 2-fold stage and these animals arrest as scrawny, starved-looking L1 that fail to show proper pharyngeal pumping but display no obvious pharyngeal morphology defects (**Fig.5E**). Taken together, the phenotypes of the *cis*-regulatory allele, as well as the neuron-specific removal of floxed *pha-4* let us conclude that *pha-4* function extends beyond embryonic patterning to a function in postmitotic neurons.

### Effect of *pha-4* removal on neuron differentiation

We used both the *cis*-regulatory allele of *pha-4* as well as the postmitotic Cre-mediated excision of the floxed *pha-4* locus to ask whether postmitotic removal of *pha-4* expression affected PEN differentiation. Analyzing *unc-17/VAChT* expression in the *cis-*regulatory allele, as well as *eat-4/VGluT* and *cat-1/VMAT* expression in the floxed *pha-4* allele, showed gene expression defects in cholinergic, glutamatergic and serotonergic PENs (**Fig.6A**). Panneuronal identity, marked by an *unc-75* reporter (Leyva-Diaz et al. 2025), is unaffected after postmitotic *pha-4* removal (**Fig.6A**), leading us to conclude that *pha-4* participates in regulating the proper identity of PENs, but not their survival.

**Figure 6:**
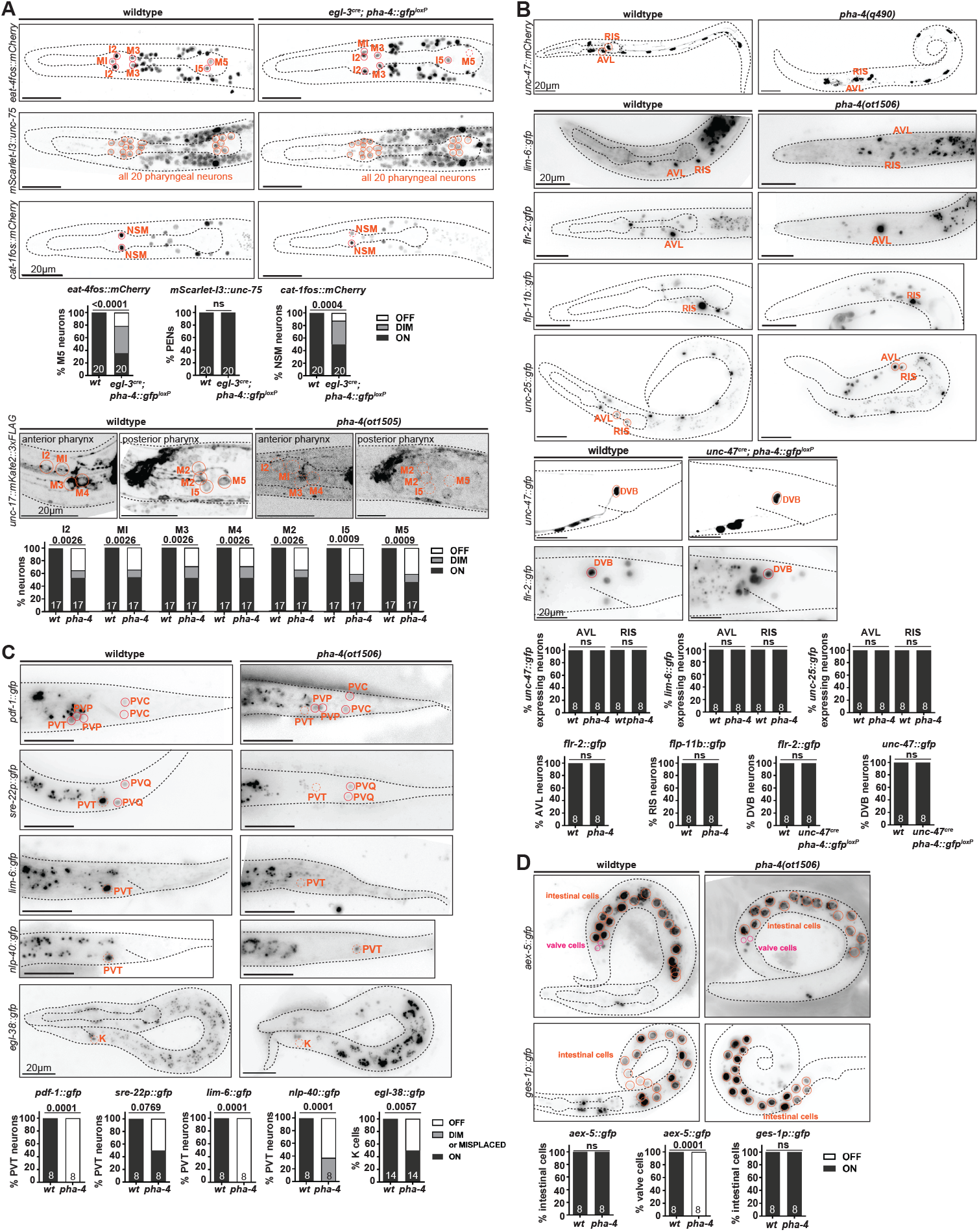
Effect of neuron-specific removal of *pha-4* on PEN & HEN differentiation. **A:** Loss of *pha-4* affects the expression of neurotransmitter identity genes, but not panneuronal identity genes. Images of L1 wildtype and L1 lethal larva show reporter transgenes *eat-4fosmid::mCherry* (*otIs518*), mScarlet-I3::*unc-75* (*ot1539*), and *cat-1fosmid::mCherry (otIs625*), in an *Cre::egl-3*(*syb5804*) *pha-4*^*loxP*^ (*ot946 ot1078*) background. Images of an L1 lethal larva show reporter allele *unc-17::mKate2::3xFLAG* (*ot907*) in the *pha-4(ot1505)* background. L1 lethal animals most likely to be homozygous *pha-4(ot1505)* were picked from a heterozygous plate based on scrawny appearance and developmental arrest, thus some heterozygous animals may have been scored, dampening the mutant phenotype quantified. Statistical analysis was performed using Fisher’s exact test. N is indicated within each bar and represents the number of animals scored. **B:** Loss of *pha-4* does not affect cell-specific identity genes in AVL, RIS, and DVB. Images of L1 lethal larva show reporter transgene *unc-47::mCherry* (*otIs348*) in the *pha-4(q490)* null background, *lim-6::gfp* (*wgIs387* reporter transgene), *flr-2::gfp* (*syb4861* reporter allele), *flp-11b::gfp* (*ynIs40* reporter transgene), and *unc-25::gfp* (*ot1372* reporter allele) in the *pha-4(ot1506)* null mutant background, and *unc-47::gfp* (*oxIs12* reporter transgene) and *flr-2::gfp* (*syb4861* reporter allele) in the *unc-47p::Cre* (*arTi479*); *pha-4*^*loxP*^(*ot1078 ot946*) background. Statistical analysis was performed using Fisher’s exact test. N is indicated within each bar and represents the number of animals scored. **C:** Loss of *pha-4* affects cell-specific identity genes in PVT and rectal K cells. Images of L1 lethal larva show reporter transgenes *pdf-1::gfp* (*syb3330* reporter allele), *sre-22::gfp* (*otEx8333* reporter transgene), *lim-6::gfp* (*wgIs387* reporter transgene), *nlp-40::gfp* (*syb3208* reporter allele), and *egl-38::gfp (wgIs171* reporter transgene) in the *pha-4(ot1506)* full locus deletion background. Statistical analysis was performed using Fisher’s exact test. Sample size is indicated within each bar and represents the number of animals scored. **D:** Loss of *pha-4* does not affect intestine-specific identity genes. Images of L1 lethal larva show reporter allele *aex-5::gfp* (*ot1532*) and *ges-1p::gfp* reporter transgene *otIs904* (Sural et al. 2025) in the *pha-4(ot1506)* null mutant background. Statistical analysis was performed using Fisher’s exact test. N is indicated within each bar and represents the number of animals scored.

To assess the impact of *pha-4* on the differentiation of the extrapharyngeal neurons that we newly identified to express *pha-4* (RIS, AVL, PVT, and DVB), we did not rely on neuron-specific removal of *pha-4*, but rather first examined a complete *pha-4* null allele, to assess the strongest possible phenotype. In spite of the overall disorganization of the head region in *pha-4* null mutants, molecular markers for RIS, AVL and PVT are selectively enough expressed to allow for proper assessment of expression in *pha-4* null mutants. For the RIS and AVL neurons, we used terminal function genes previously shown to be critical for the function of these neurons, the vesicular transporter *unc-47/VGAT* (AVL and RIS), the GABA biosynthetic enzyme *unc-25/GAD*, and the neuropeptides *flp-11* (RIS) and *flr-2* (AVL). Expression of these markers is not affected in L1-arrested *pha-4(ot1506)* null mutant animals (**Fig.6B**). Notably, *flr-2* is normally also expressed in PENs (Vidal et al. 2022) and we find PEN expression to be lost in *pha-4* null mutants (**Fig.6B**).

To assess the effect of neuron-specific *pha-4* removal on DVB, which is born after the L1 stage (i.e. after *pha-4* null mutant animals die), we removed *pha-4* cell-specifically from DVB by driving Cre recombinase under control of the GABA neuron-specific promoter of the *unc-47/VGAT* locus. An *unc-47prom::Cre* miniMOS line indeed removed endogenously *gfp-*tagged *pha-4* from the RIS, AVL and DVB neurons, but not the PVT neurons, as expected (**Fig.7A**). Post-embryonic loss of *pha-4* from DVB does not affect the expression of molecular markers *flr-2*, and *unc-47/VGAT* (**Fig.6B**). However, we cannot exclude the possibility that the Cre-mediated gene removal is too late to affect differentiation of the neurons. Nevertheless, considering the result with the null allele, it appears that in contrast to strong differentiation defects observed in the PENs, the RIS, AVL, and DVB neurons are generated and appear to adopt their proper fate in the absence of *pha-4* function.

**Figure 7:**
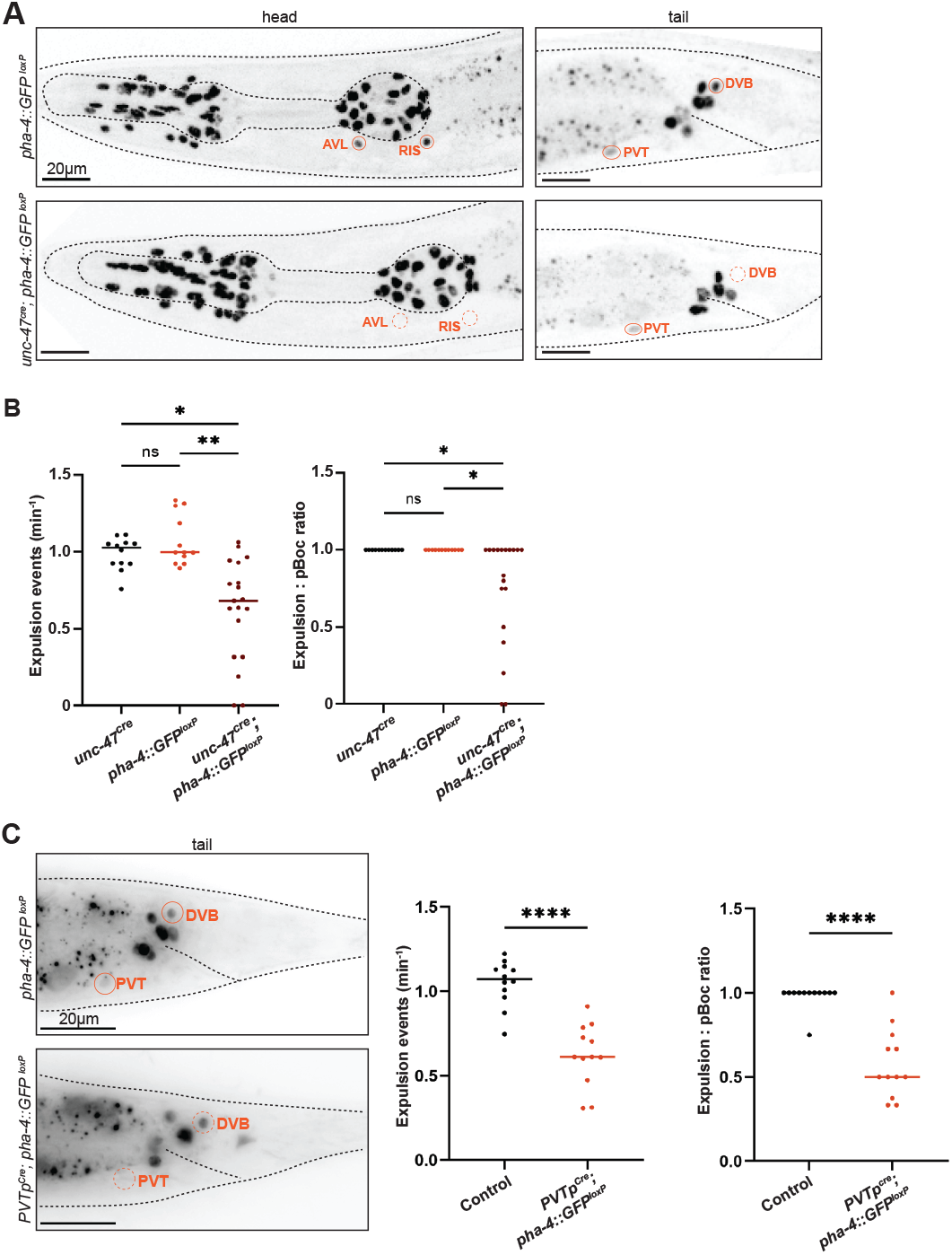
*pha-4* acts in HENs to control defecation behavior. **A:** Cell-specific removal of *pha-4* from the *gfp-*tagged, floxed *pha-4* locus (*ot946 ot1078*), generated via the CRISPR/Cas9 system, via *unc-47p::Cre* (*arTi479*). Elimintion of PHA-4::GFP expression was confirmed at the day 1 adults stage. **B:** *pha-4* removal from AVL, RIS, and DVB affects defecation behaviour. Frequency of expulsion and expulsion:pBoc ratio in adult worms with cell-specific removal of *pha-4* locus (floxed reporter allele *ot946 ot1078*) via *unc-47p::Cre* (*arTi479*). **C:** Left panel: neuron-specific removal of PHA-4::GFP (floxed reporter allele *ot946 ot1708*) via *sre-22* promoter, PVT*p::Cre* (*otEx343*) was confirmed in adult animals. All remaining PHA-4::GFP cells in the tail region are non-neuronal PHA-4(+) cells. Right panel: *pha-4* removal from PVT affects defecation behavior. Frequency of expulsion in adult worms with PVT-specific removal of *pha-4* (floxed reporter allele *ot946 ot1078*) via PVT*p::Cre* (*otEx343*). Data points representing each individual worm assayed are plotted, and horizontal line represents median value of biological replicates in (B-C). *, **, **** and ns represent P < 0.05, P < 0.01, P < 0.0001 and not significant, respectively, in Dunn’s multiple comparison test after Kruskal-Wallis test (B), and Mann-Whitney test (C).

In the PVT neurons, *pha-4* appears to play an earlier patterning role. PVT is the sister of rectal cells that Mango and colleagues had noted to be absent in *pha-4* mutants (Mango et al. 1994). Consistent with a lineage defect, we find that in *pha-4* null mutants, expression of a terminal marker for PVT identity, *sre-22*, is frequently lost, as is expression of a reporter allele of the *pdf-1* neuropeptide (**Fig.6C**). The *nlp-40* neuropeptide reporter allele shows reduced expression in a tail cell outside of the pre-anal ganglion region, where PVT normally resides (**Fig.6C**). We further corroborated a potential lack of differentiated PVT by examining the expression of a regulator of PVT identity, the LIM homeobox gene, *lim-6* (Hobert et al. 1999; Aurelio et al. 2003). We find no observable *lim-6* signal in the tail region (**Fig.6C**), consistent with a loss of the PVT neuron.

We also examined the impact of *pha-4* on the rectal epithelial K cell, which is lineally related to PVT (**Fig.1B**), and the precursor of the postembryonically born DVB hindgut innervation neurons. We found that expression of the K cell marker, *egl-38*, is affected in *pha-4* null mutants (**Fig.6C**), again consistent with *pha-4* having early developmental defects in generating components of the hindgut. This is further supported by our analysis of the expression of *aex-5*, which codes for an endopeptidase involved in defecation behavior (Mahoney et al. 2008). An *aex-5* reporter allele that we generated by CRISPR/Cas9 genome engineering (Sural et al. 2025) is expressed in the rectal gland cells (one of which, rectD, also lineally related to PVT; **Fig.1B**) and in intestinal-rectal valve cells (virL/R) and this expression is lost in *pha-4* null mutants (**Fig.6D**).

*lim-6* is also a terminal selector of AVL and RIS neuron identity and, again consistent with the lack of differentiation defects of AVL and RIS (Hobert et al. 1999; Gendrel et al. 2016), we find *lim-6* expression in these neurons to be apparently unaffected in *pha-4* null mutants (**Fig.6B**). We also noted ectopic *lim-6* expression in the head, perhaps indicating that pharyngeal cells that fail to differentiate properly in *pha-4* mutants undergo a cell fate switch (**Fig.6B**).

### *pha-4* acts in GABAergic HENs and in PVT to control defecation behavior

We found that in *pha-4(ot1506)* null mutants, the expression of several markers of terminal differentiation of intestinal cells (*ges-1* and *aex-5)* is unaffected (**Fig.6D**). Previous work has shown that *pha-4* is required for proper function and homeostasis of mature intestinal cells (Panowski et al. 2007; Wu et al. 2018; Torzone et al. 2023), indicating that in specific cellular contexts, *pha-4* may not have a role in the proper differentiation of a cell type, but rather in its proper functioning. We set out to test whether a similar scenario applies to the extrapharyngeal neurons (HENs, RIS and PVT), whose overal differentiation appears unaffected in *pha-4* null mutants. To avoid the larval lethality associated with either *pha-4* null mutants or with panneuronal removal of *pha-4*, we set out to remove *pha-4* selectively from HENs (AVL, DVB). To this end, we expressed Cre recombinase specifically in GABAergic neurons using the *unc-47/VGAT*promoter. An *unc-47prom::Cre* miniMOS line indeed removed endogenously *gfp-*tagged *pha-4* from the RIS, AVL and DVB neurons, but not from PVT or other rectal cells, conforming with the specificity of the *unc-47prom* driver (**Fig.7A**). Since AVL and DVB neurons, both motor neurons that innervate enteric muscle, were previously shown to control defecation behavior (McIntire et al. 1993; Choi et al. 2021), we analyzed this behavior in animals lacking *pha-4* in GABAergic neurons and indeed observed defecation defects. While the rate of pBoc events in these worms was only mildly affected, not all pBoc events were followed by expulsion events (**Fig.7B**). In some worms, expulsion events were entirely absent for the six observed defecation cycles **(Fig.7B)**, which is consistent with the phenotype of worms in which AVL and DVB are ablated (McIntire et al. 1993; Choi et al. 2021). Removing *pha-4* from GABAergic neurons does not affect pharyngeal pumping behavior (**Fig.7B; Fig.S3**).

To assess whether PVT requires *pha-4* to regulate defecation behavior, we again employed the above-described *sre-22prom* PVT driver to express Cre in the PVT neuron of animals that contain a floxed copy of the *pha-4* locus. These animals also exhibit significant defects in the expulsion step of defecation, with only about half of pBocs being followed by an expulsion step (**Fig.7C**). These defects were similar to those observed upon selective removal of *egl-3* from PVT (**Fig.3**). We conclude that *pha-4* is required for features of PVT that are critical for its function in controlling defecation behavior.

### *pha-4* is continuously required to sustain function of HENs and PENs

While Cre-mediated removal of *pha-4* establishes a function for *pha-4* in postmitotic PENs and HENs, it does not address whether *pha-4* may also function post-embryonically to maintain function of these neurons through to adulthood. Previous work using a temperature-sensitive allele of the *pha-4* locus indicated that *pha-4* removal at the first larval stage resulted in animal lethality (Gaudet and Mango 2002), but it was not clear in which tissue type *pha-4* was required, nor whether this requirement persisted into later stages of animal life. To address this issue, we used the auxin-inducible degron system (Zhang et al. 2015). We tagged the endogenous *pha-4* locus with an auxin-inducible degron (AID) sequence (**Fig.8A**) and generated a transgenic line in which the TIR1(F79G) ligase (Hills-Muckey et al. 2022) is driven by the *pha-4prom2* driver described above, which is expressed in the PENs, HENs (AVL and DVB) and RIS, but not in PVT. After addition of the auxin-derivative 5-Ph-IAA, we observe that GFP::loxP::3xFLAG::AID2-tagged PHA-4 is indeed depleted from the PENs, AVL and RIS, but not PVT or DVB (**Fig.8B**). Using this system, we observed feeding and defecation behavior of adult animals that have either been treated since embryonic, L1 or L4 larval stages with 5-Ph-IAA (**Fig.8C**). We conclude that PHA-4 is required continuously after embryonic development for maintainance of enteric functions.

**Figure 8:**
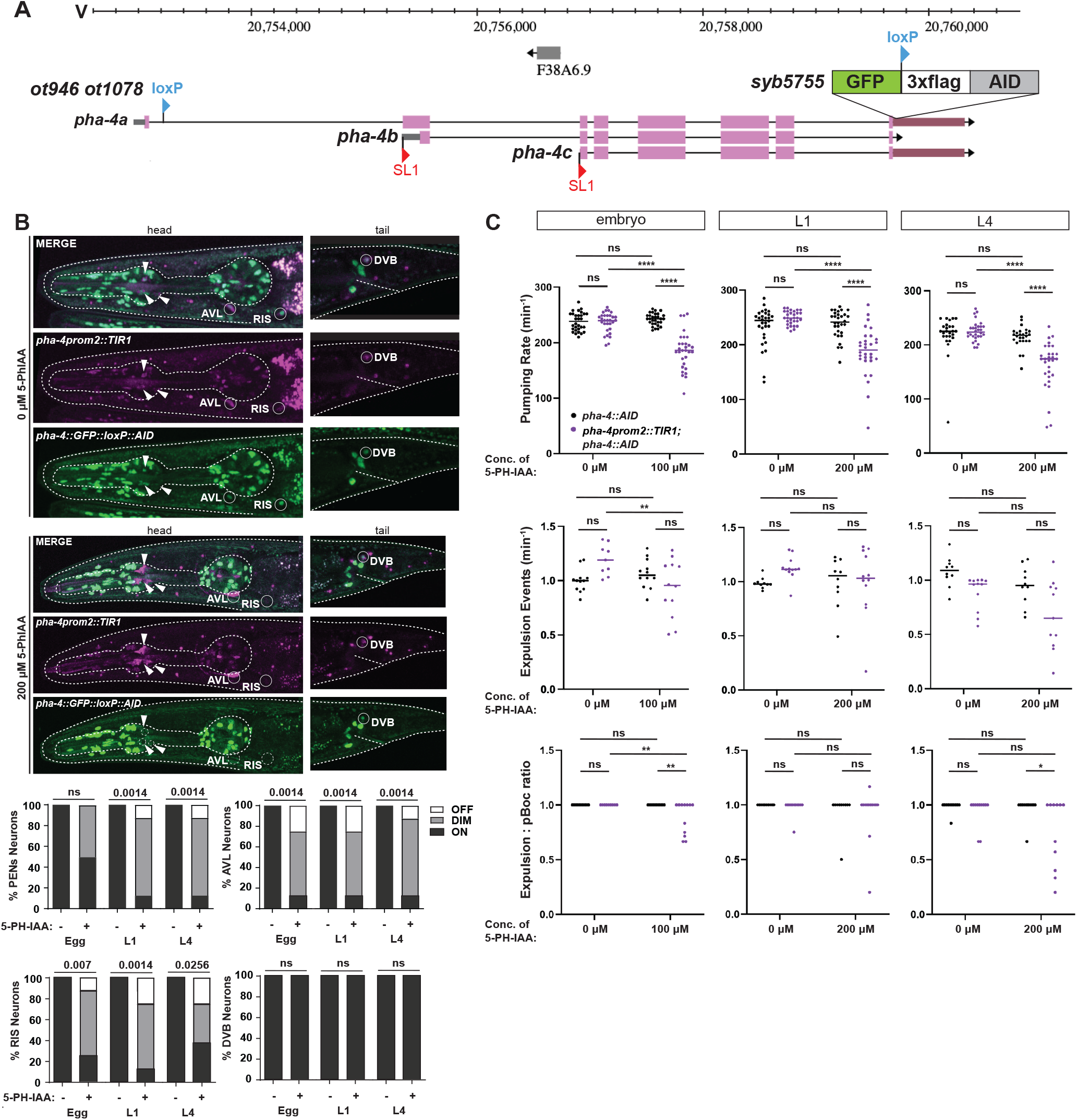
*pha-4* is continuously required in enteric neurons to maintain enteric function. **A**: *pha-4* locus schematic showing location of an additional AID tag inserted via CRISPR/Cas9 engineering into the already *gfp*-tagged *pha-4* reporter allele *ot1078 ot946*, resulting in *pha-4*(*ot946 ot1078 syb5755*), which we label as *pha::gfp::AID* in the ensuing panels. **B**: Temporally controlled removal of PHA-4 in *pha-4*(*ot946 ot1078 syb5755)* animals from enteric neurons via pha-4prom2::*TIR1(F79G)*(*otIs908*) in the presence of 5-Ph-IAA. Animals were treated with either ethanol only (0 μM 5-Ph-IAA) or 200 μM 5-Ph-IAA, starting at various stages, and GFP expression in *pha-4(+)* neurons was characterized. White arrows point to *TIR1(F79G)* and *pha-4*(*ot946 ot1078 syb5755)* expression in pharyngeal neurons. Statistical analysis was performed using Fisher’s exact test. N is indicated within each bar and represents the number of neurons scored. **C:** Temporally controlled removal of *pha-4* from enteric neurons results in defects in enteric behaviors. Pharyngeal pumping, frequency of expulsion, and expulsion:pboc ratio in adult worms treated with auxin starting at different developmental stages. Animals were treated with either ethanol only (0 μM 5-Ph-IAA) or indicated concentrations of 5-Ph-IAA. Data points representing each individual worm assayed are plotted, and the horizontal line represents the median value of the biological replicates in (C-D). *, **, **** and ns represent P < 0.05, P < 0.01, P < 0.0001 and not significant, respectively, in Sidak’s multiple comparisons test after two-way ANOVA (C), and a Mann-Whitney test (D).

## DISCUSSION

The role of FoxA genes in gut development is evident across animal phylogeny. Previous studies on the *C. elegans* FoxA ortholog PHA-4 have firmly established its role as an organ selector gene for foregut development (Mango et al. 1994; Mango 2009). Target genes for PHA-4 have been defined during this embryonic patterning function in different parts of the foregut (Gaudet and Mango 2002; Gaudet et al. 2004; Fakhouri et al. 2010) and, on a mechanistic level, PHA-4 has been shown to act as a pioneer factor to enable gene activation (Hsu et al. 2015). We have extended here the description of PHA-4 function by revealing postmitotic functions of PHA-4 during terminal differentiation of enteric neurons, not only of the foregut, but also the hindgut, as well as other neurons that control the function of the fore- or hindgut. These postmitotic functions extend beyond into larval and adult stages. Since pioneer factors – of which the vertebrate PHA-4 ortholog FoxA is the paradigmatic example - are generally thought to provide the initial trigger for opening chromatin (Zaret and Carroll 2011), a continuous requirement for PHA-4 indicates functions of PHA-4 that go beyond such pioneer function. We propose that continuously expressed PHA-4 cooperates within the foregut enteric nervous system with a previously identified pan-enteric homeodomain transcription factor, the Six-type homeodomain protein CEH-34, that is also required to initiate and maintain enteric neuron differentiation (Vidal et al. 2022). A direct role of PHA-4 in maintaining, together with CEH-34, the fully differentiated state of pharyngeal enteric neurons is supported by the presence of ChIP-Seq peaks for PHA-4 (identified by the ModEncode project; (Gerstein et al. 2010)) in the vicinity of terminal marker genes, including, for example, the *cat-1/VMAT* locus whose expression we find to be affected by postdevelopmental removal of PHA-4.

Perhaps most remarkably, our discovery of expression of PHA-4 in extrapharyngeal neurons points to a fundamental relatedness of HENs and PENs, but also identified other neurons that control the enteric system. PHA-4-expressing AVL and DVB neurons have long been recognized as hindgut innervating neurons that control defecation behavior. The extrapharyngeal RIS interneuron, that we discover here to also express PHA-4, is a known regulator of pharyngeal pumping (Steuer Costa et al. 2019). Even though its cell body is located outside the foregut in the ventral ganglion on its head, the extension of its unipolar process along the nerve ring that wraps around the isthmus of the pharynx, puts RIS in a suitable position to control pharyngeal pumping apparently via the release of neuropeptides (FLP-11) perceived by neuropeptide receptors expressed in pharyngeal enteric neurons. The expression of PHA-4 in the PVT neuron guided us to explore a potential function of PVT in enteric behavior as well. We discovered that PVT plays a role in controlling the expulsion step of defecation, apparently mediated via neuropeptide release. Removing *pha-4* from PVT using a cell-specific Cre driver recapitulated the expulsion defects seen in worms where PVT was silenced, suggesting that *pha-4* is required for the enteric function of PVT.

Previous work has suggested that the neuropeptide NLP-40 is secreted from the intestine to stimulate proper levels of GABA release in AVL and DVB (Wang et al. 2013). Since we found *nlp-40* to also be very strongly expressed in PVT and since the NLP-40 receptor AEX-2 is expressed not only in AVL and DVB, but also in PVT (Ripoll-Sanchez et al. 2023), it is conceivable that PVT may operate as an amplifier. It may detect NLP-40 released from the intestine via the AEX-2 receptor to promote further NLP-40 release from PVT to stimulate AEX-2 in AVL and DVB during the expulsion step.

With its extension of a long process into the nerve ring from the tail of the animal, PVT represents a potential route of communication from the brain to the hindgut, akin to the vagus nerve. The lineage relationship of PVT with other parts of the hindgut (**Fig.1B**) could have already predicted a function of PVT in controlling hindgut function. Yet, we note that lineage relationships of functionally related neurons are certainly not a universal feature, within the gut (e.g. the hindgut innervating neuron AVL has no lineage relationship to any part of the gut) or outside the gut (e.g. amphid sensory neurons have very limited lineal relationships). Lastly, we note that the function of the peptidergic RIS and PVT neurons in controlling enteric function can be viewed as conceptually similar to the control of vertebrate enteric nervous system function (e.g. gut motility) by neuropeptides produced by the CNS (Furness 2012).

While PHA-4 may act together with (and upstream of) a terminal selector, CEH-34, to control multiple aspects of PENs in a master regulatory manner, PHA-4 function in extrapharyngeal neurons (AVL, DVB, RIS, PVT) may be restricted to controlling select aspects of the functional properties of these neurons. Notably, all these four neurons utilize a shared LIM homeodomain transcription factor for their proper differentiation, *lim-6* (Hobert et al. 1999; Aurelio et al. 2003; Gendrel et al. 2016). In each of these neurons, PHA-4 may assist LIM-6 in controlling some critical aspect of their function. A similarly selective mode of action of PHA-4 in controlling cell function rather than cell differentiation is observed in the midgut endoderm, which also forms properly in *pha-4* mutants, but which displays a number of functional defects, such as in lipid homeostasis (Wu et al. 2018; Torzone et al. 2023). Hence, our findings demonstrate that PHA-4 acts in a highly contextualized environment. In some cells it acts to control cell differentiation in a “master regulatory manner”, in combination with other transcription factors, while in yet other cell types its function is committed to controlling a highly selective subset of as yet undiscovered genes to control select aspect of their functional properties.

We conclude that PHA-4 is currently the only gene whose expression is tightly restricted to every single neuron of an enteric nervous system, where it fulfills a number of diverse functions from pioneer function role in early patterning, to the initiation of terminal differentiation programs to the maintenance of enteric neuron functionality throughout the life of the animal.

## MATERIAL AND METHODS

### Strains

All *C. elegans* strains were cultured at 20°C on nematode growth medium (NGM) plates seeded with *Escherichia coli* (OP50 strain) bacteria as a food source unless stated otherwise. The wildtype strain used is Bristol N2. All experiments were performed on hermaphrodites. A comprehensive list of the strains utilized in this study is available in **Table S1**. Details for the generation of several transgenic and CRISPR/Cas9-engineered strains are provided in the Supplemental Material and Methods section.

### Reagents for PVT neuron visualization and manipulation

To identify drivers for PVT neuron expression, pZW28 [*cnc-11p::gfp::h2b::tbb-2 3’UTR*] was generated by cloning a 757 bp *cnc-11* promoter fragment (-757 to - 1 position with respect to the start codon of *cnc-11*). pZW30 [*sre-22p::gfp::h2b::tbb-2 3’UTR*] and pZW32 [*sre-22p:: HisCl::sl2::gfp::tbb-2 3’ UTR*] were generated by cloning a 1,144 bp *sre-22* promoter fragment (-1,144 to -1 position with respect to the start codon of *sre-22*). The *tbb-2* 3’ UTR sequence is a 156 bp sequence immediately downstream of the stop codon of the *tbb-2* gene. The *sre-22* driver was preferred for further usage (HisCl and Cre expression) over the *cnc-11* driver because of its greater specificity, with the only non-PVT site of expression being dim expression in the PVQ neurons. While we cannot exclude a role for PVQ in interpretating the results of the *sre-22prom*-driven HisCl silencing, the *sre-22prom* driven excision of *pha-4* from PVT, with ensuing defects matching those of *sre-22prom::HisCl*, argues against a contribution from PVQ, since *pha-4* is not expressed in PVQ.

All plasmids used in this study were generated via NEBuilder HiFi DNA assembly using the manufacturer’s protocol. All plasmid sequences were validated using Sanger (Azenta) and/or Oxford Nanopore (Plasmidsaurus) sequencing. Prior to microinjection, all plasmids were linearized using a restriction site in the plasmid backbone. The linearized plasmids were then injected into both gonadal arms of young adult animals. F1 progeny were selected based on the expression of the co-injection marker, and those that transmitted the array to F2 progeny were used to establish transgenic lines. Three independent lines (derived from independently injected P0 animals) for each injection type were utilized in subsequent experiments.

*nlp-40* and *pdf-1* expression in PVT, intially suggested by the CeNGEN atlas (Taylor et al. 2021), was examined by inserting an T2A::3xNLS::GFP cassette at the 3’ end of the *nlp-40* and *pdf-1* coding sequences (SunyBiotech). This reporter cassette was subsequently shown to produce lower reporter expression than other reporter cassette (Sun et al. 2023), which explains why the *nlp-40* reporter allele shows expression in PVT, but not the midgut, as previously reported (Wang et al. 2013).

### Assays for larval arrest of *pha-4* mutants

To assess the consequences of neuronal removal of *pha-4*, we analyzed the progengy of animals in which either the neuronal enhancer of the *pha-4* locus was deleted (*ot1505 ot946* allele) or in which floxed *pha-4* was removed with a panneuronal Cre driver. For each parental genotype, ten age-synchronized L4 stage hermaphrodites were transferred from uncrowded, non-starved plates to ten individual fresh NGM plates seeded with *E. coli* OP50 bacteria. After 24 hours at 20°C, parent worms were transferred to new NGM plates seeded with OP50 bacteria. The number of live larvae and unhatched embryos was counted for each plate after an additional 24 hours at 20°C. Parent worms were iteratively transferred and progeny counted in 24 hour intervals two additional times, and the total hatch rate was pooled for each individual parent.

### Measurement of enteric behaviors

All pharyngeal pumping assays were performed on freely moving animals. Age-synchronized adult worms were transferred from uncrowded, non-starved plates to fresh NGM plates seeded with a uniform thin layer of *E. coli* OP50 bacteria. After a 5-minute acclimatization period, worms were placed under a Nikon Eclipse E400 upright microscope equipped with differential interference contrast (DIC) optics. Before recording behavior, worms were allowed to acclimate to the light source intensity under the microscope for an additional 5 minutes. The movement of the grinder of the pharynx was then recorded with a hand-held tally counter using a 20x air objective lens. The number of grinder movements in 20 seconds was recorded and multiplied by three to obtain pharyngeal pumps per minute. The pharyngeal pumping rate was recorded from at least 8 animals each day and data was pooled from at least on two independent days.

All defecation assays were performed on freely moving animals. Age-synchronized adult worms were transferred from uncrowded, non-starved plates to fresh NGM plates seeded with a uniform thin layer of *E. coli* OP50 bacteria. After a 5-minute acclimatization period, worms were placed under a Nikon Eclipse E400 upright microscope equipped with DIC optics. Before recording behavior, worms were allowed to acclimate to the light source intensity under the microscope for an additional 5 minutes. Defecation behavior was observed under the 20x air objective lens. Each animal was observed for a 6 min period, during which the timings of contraction of posterior body muscles (pBoc) and expulsion (Exp) muscle contraction that releases food contents from the anal opening were recorded. Defecation behavior was recorded from at least six animals on each day and data was pooled from at least two independent days.

### Histamine-mediated chemogenetic silencing of neurons

The inducible silencing of RIS and PVT neurons was conducted using the histamine-gated chloride channel HisCl1 from *Drosophila melanogaster* and by the addition of exogenous histamine as previously described (Pokala et al. 2014). Histamine dihydrochloride (Sigma-Aldrich H7250) was dissolved in water to prepare 1M stock solutions. NGM Agar containing 0 mM (control), 8 mM, and 10 mM histamine dihydrochloride was added to 60mm plates. Age-synchronized adults were transferred to NGM agar control and histamine plates immediately prior to experiments. After a recovery period, worms were observed for pharyngeal pumping and defecation behavior under a Nikon Eclipse E400 upright microscope equipped with DIC optics. For L1 molting experiments, worms were age-synchronized in the L1 stage and observed for pharyngeal pumping and defecation behavior 9 hours after being added to NGM control and histamine plates at 23°C.

### Auxin-mediated protein degradation

Auxin-inducible degron (AID)-tagged proteins are conditionally degraded when exposed to 5-Ph-IAA in the presence of TIR1(F79G). All AID2 experiments were carried out as described previously with some modifications (Sural et al. 2024). To generate the experimental strain, the conditional allele *pha-4(ot946 ot1078 syb5755)* was crossed with *otIs908* which expresses TIR1(F79G) in all neurons of the pharyngeal nervous system and AVL, RIS, and DVB. The synthetic auxin analog 5-Ph-IAA was acquired from BioAcademia (#30-003-10) and dissolved in ethanol (EtOH) to prepare 100 mM stock solutions. NGM agar plates seeded with a uniform lawn of *E. coli* OP50 bacteria were coated with 100 μL of freshly prepared 5-Ph-IAA stock solution and allowed to dry and diffuse overnight at room temperature to final concentrations of 100 μM and 200 μM. Age-synchronized embryos (obtained via the alkaline bleaching method), L1s, or L4s were transferred to the plates one day after adding 5-Ph-IAA, and the animals were grown at 24°C for an additional 24 hours until they reached the desired developmental stage for the experiment. In control conditions, worms were transferred onto EtOH-coated plates. All plates were shielded from light for the duration of the experiment to limit degradation of 5-Ph-IAA.

### Microscopy

Worms were anesthetized using 100 mM sodium azide (NaN_3_) and mounted on 5% agarose pads on glass slides. Animals were imaged on an inverted Zeiss LSM 980 laser scanning confocal microscope using a 40x water immersion objective lens, or a Zeiss Imager Z2 upright microscope equipped with Colibri 7 LEDs (Zeiss) using a 40x oil immersion objective lens. For each image, Z-stacks of uniform 0.31 μm - 1 μm thickness (depending on age of the worm) were obtained for the entire sample.

Reporter gene expression in different neurons was visualized in wildtype and mutant animals and usually assigned to one of the following categories: ‘on’ (fluorescence levels comparable to wild-type animals), ‘dim’ (fluorescence still detectable but much dimmer than wildtype animals), or ‘off’ (fluorescence not detectable) by the observer.

### Statistical analysis

All images were processed and analyzed using ImageJ FIJI (Schindelin et al. 2012). All plotting and statistical tests were performed on GraphPad Prism 10. Sample sizes for pharyngeal pumping and defecation assays are a minimum of 32 and 12 animals respectively, based on standards established in prior studies using these behavioral readouts. These sample sizes have been shown to reliably detect physiologically relevant differences. For microscopy images, a minimum of 8 animals were imaged per condition, and a representative image is shown. The figure legends include the statistical tests applied in each figure along with the corresponding *P* values. *P* < 0.05 was considered to be statistically significant.

## ACKNOWLEDGEMENT

We thank Qi Chen for expert help in generating transgenic strains, and Rudy LeDuc, Emma Grungold and Tristian Wiles for their involvement in generating the *unc-47prom::cre* line and David Raizen for providing strains. Some strains were provided by the CGC, which is funded by NIH Office of Research Infrastructure Programs (P40 OD010440). This work was funded, in part, by the NIH (R01 NS039996) and the Howard Hughes Medical Institute.

## AUTHOR CONTRIBUTIONS

Conceptualization: OH

Methodology: ZW, WXC, ELD, MR, SS, MAA

Investigation: ZW, WXC, ELD, MR, SS, MAA

Visualization: ZW, WXC, SS

Supervision: OH

Writing—original draft: OH

Writing—review & editing: ZW, WXC, ELD, MR, SS, MAA, OH

