## Supplementary Information for "PHA-4/FoxA controls the function of pharyngeal and extrapharyngeal enteric neurons of *C. elegans*"

### Supplemental Material and Methods

#### Transgenic line generation

Two *pha-4* promoter fragments were used to drive expression of a GFP reporter. pWXC065 [*pha-4prom1::gfp::unc-54 3'UTR*] was generated by cloning a 1963 bp promoter fragment upstream of the longest isoform of pha-4 (-1954 to +9 position relative to the start codon of *pha-4a*). pWXC067 [*pha-4prom2::gfp::unc-54 3'UTR*] was generated by cloning a 2393 bp fragment including nearly the entire first intron of *pha-4a* isoform and up to the second codon of the *pha-4b* isoform (position +53 to +2445 relative to the start of *pha-4a*). These promoter fragments were cloned into the multiple cloning site of the *C. elegans* GFP expression vector pPD95.75 (Fire Vector Kit). Transgenic worms carrying extrachromosomal arrays of these reporter constructs were generated by injecting these plasmids into a *pha-1(ts) allele* alongsite a *pha-1(+)* rescue plasmid, and array-carrying worms were selected for at 25°C, resulting in line *otEx8085*. The *pha-4prom2::gfp* plasmid was also independently integrated into the genome (resulting in the *otIs946* transgene), using the Fluorescent Landmark Interference (FLInt) technique (Malaiwong et al. 2023). We used slightly modified concentrations compared to the previous protocol. Specifically, to prepare the FLInt injection mixes, tracrRNA (final concentration of 100 ng/μL) and a crRNA (final concentration: 56 ng/μL) targeting the tdTomato locus were combined and incubated at 95°C for 5 minutes, followed by a 5-minute incubation at 10°C. Cas9 (final concentration: 250 ng/μL) was then added, mixed by pipetting, and incubated at 25°C for 10 minutes. Afterward, all plasmids were added at specified concentrations. The final volume was brought to 20 μL using nuclease-free water, followed by centrifugation at 17,900 xg for 2 minutes. The supernatant was then used for microinjections into *oxTi553[eft-3p::tdTomato::H2B::unc-54 3' UTR + Cbr-unc-119(+)] V* animals. Injected animals were maintained at room temperature, and F1 progeny expressing the co-injection marker were singled onto separate plates. F2 progeny were selected from F1 plates where at least 75% of the animals expressed the co-injection marker. F2 plates that showed transmission of the co-injection marker in 100% of the progeny were used to establish lines.

The *pha-4prom2* fragment was also used to drive additional reagents, namely TIR1(F79G) for auxin-dependent protein degradation (*otIs908*) and GCaMP6s for visualizing activity of enteric neurons (*otIs898*). The *otIs908* strain is described in (Sural et al. 2025). To drive GCaMP expression with *pha-4prom2* fragment, pWXC072 [*pha-4prom2::3xNLS::ceGCaMP6s::unc-54 3' UTR*] was cloned by swapping out the promoter region of previously published pEY13 [*arrd-4p::3xNLS::ceGCaMP6s::unc-54 3'UTR*] (Yemini et al. 2021), containing the *C. elegans* codon-optimized GCaMP6s sequence. The *arrd-4* promoter was excised by restriction digest by SphI and XmaI, and PCR-amplified *pha-4prom2* promoter fragment was inserted using the HiFi Assembly kit (NEB). Plasmid DNA sequence was validated by Sanger sequencing (Azenta). This plasmid was integrated via FLInt-mediated integration (as described above), resulting in the *otIs898* transgene.

We generated Cre driver lines in different ways. PVT*prom::Cre* (*sre-22p::2xFlag::NLS::Cre::tbb-2 3'UTR, unc-122::gfp*) was generated by amplifying the *2xFlag::NLS::Cre* region from pMA193. The amplified *2xFlag::NLS::Cre* regulatory region was cloned into the pZW30 backbone amplified from the *tbb-2* 3'UTR to the 1,144 bp *sre-22* regulatory region to generate pZW37 *sre-22p::2xFlag::NLS::Cre::tbb-2 3'UTR*. Before microinjection, pZW37 was linearized using a single cutter restriction enzyme that cuts in the plasmid backbone. Linearized pZW37 was injected at a concentration of 2 ng/uL into both gonadal arms of young adult animals. F1 progeny was picked based on expression of the co-injection marker *unc-122p::gfp* (5 ng/uL) and animals that transmitted the array to F2 progeny were used to generate transgenic lines. Two independent lines (obtained from independent injected P0 animals) were used in subsequent experiments.

*Cre::ceh-48(syb5859)* and *Cre::egl-3(syb5804)* alleles contain an insertion of a *FLAG::NLS::Cre::SL2* cassette before the start codon (generated by SunyBiotech). The SL2 trans-splice site was used to ensure that both *Cre* and *ceh-48 (*or *egl-3)* were co-transcribed in the same cells. The 300bp *unc-47* regulatory region was amplified from pMG071 with SpeI and XmaI sites appended. The amplified and cut *unc-47* regulatory region was cloned into a miniMOS vector containing a version of Cre optimized for efficiency in *C. elegans* (Ruijtenberg and van den Heuvel 2015) and a 3' UTR from *unc-54*. Single-copy insertions were generated using the standard miniMOS protocol (Frokjaer-Jensen et al. 2014).

**Genetic manipulations of the *pha-4* locus**

CRISPR/Cas9 genome editing was carried out to manipulate the *pha-4* locus, using a modified version of a previously established protocol (Eroglu et al. 2023). The injection mixture for all CRISPR experiments consisted of 250 ng/µL of *S. pyogenes* Cas9 nuclease (IDT), 100 ng/µL of tracrRNA (IDT), 56 ng/µL (combined) of all crRNAs (IDT), and 100 ng/µL of each single-stranded oligodeoxynucleotide (ssODN) repair template (IDT). tracrRNA and all crRNAs were heated at 95°C for 5 minutes, followed by cooling to 10°C for 5 minutes. Cas9 was then added to the mix, thoroughly pipetted, and incubated at 25°C for 10 minutes. Next, ssODN and plasmids were incorporated into the mix, and the total volume was brought up to 20 µL with nuclease-free water. The mixture was centrifuged at 17,900 xg for 2 minutes, and the supernatant was collected for microinjection. A *dpy-10* co-CRISPR strategy was used, employing a crRNA (GCTACCATAGGCACCACGAGGTTTTAGAGCTATGCT) and ssODN (CACTTGAACTTCAATACGGCAAGATGAGAATGACTGGAAACCGTACCGCATGCGGTGCCTATGGTAGCGGAGCTTCACATGGCTTCAGACCAACAGCCTAT). F1 animals were selected based on the roller phenotype, and genotyping was performed by PCR amplification of the edited locus, followed by sequencing via Sanger (Azenta) or Oxford Nanopore (Plasmidsaurus). From confirmed heterozygous F1 plates, non-roller progeny were singled, propagated, and genotyped by PCR to establish homozygous lines.

*pha-4(ot946)* and *pha-4(ot946 ot1078)* were generated sequentially using reagents from IDT, and as previously described (Dokshin et al. 2018). The *pha-4* locus was tagged with *gfp::loxP::3xFLAG* at its 3' end to generate *pha-4(ot946)*. One crRNA (attggagatttataggttgg) and an asymmetric double-stranded gfp::loxP::3xFLAG cassette were used to insert the fluorescent tag at the C-terminal. *pha-4(ot946 ot1078)* was generated by inserting a second loxP site after *pha-4* first exon (position +278) into the existing *pha-4(ot946)* allele using the crRNA (acataagtgctcattaacag) and a ssODN donor (ccttctgtgtataatagcacataagtgctcattaaATAACTTCGTATAGCATACATTATACGAAGTTATcaggggatactcgatgtcaatatgacccggccagg). This strain was further modified (SunyBiotech) through the insertion of a GAS flexible linker and the 45 amino acid long Auxin-Inducible Degron (AID) at the end of the *gfp-loxP::3xFLAG* cassette located at the C-terminus, just before the stop codon to generate *pha-4(ot946 ot1078 syb5755)*.

Mutant alleles for *pha-4* (schematized in **Fig.5A**) were generated using *pha-4(ot946 ot1078)* as the starting strain and two crRNAs and an ssODN donor. *pha-4(ot1505 ot946)* removes a *cis*-regulatory element but leaves the protein tagged. *pha-4(ot1506)* is a full locus deletion of the coding sequence (and the reporter cassette). The sequences are as follows:

*pha-4(ot1505)* crRNAs: ACATTATTGATATAAGGGTA and AATTAAACTATTTTGTTTGA
ssODN:CGCCATCCAGTGATGAGGACATTATTGATATAAGGGTCATCAAAGAGGAGCCAGAGTCGGAACCTGACTC

*pha-4(ot1506)* crRNAs: GTCCACTGCGTATTTTTTGA and TAAAAGGAGAACGTGTGAAT
ssODN:ACCTTATACACCCTATCTGTCCACTGCGTATTTTTCACACGTTCTCCTTTTATTGGAACAGGGCATAGGA.

### Supplementary Figures

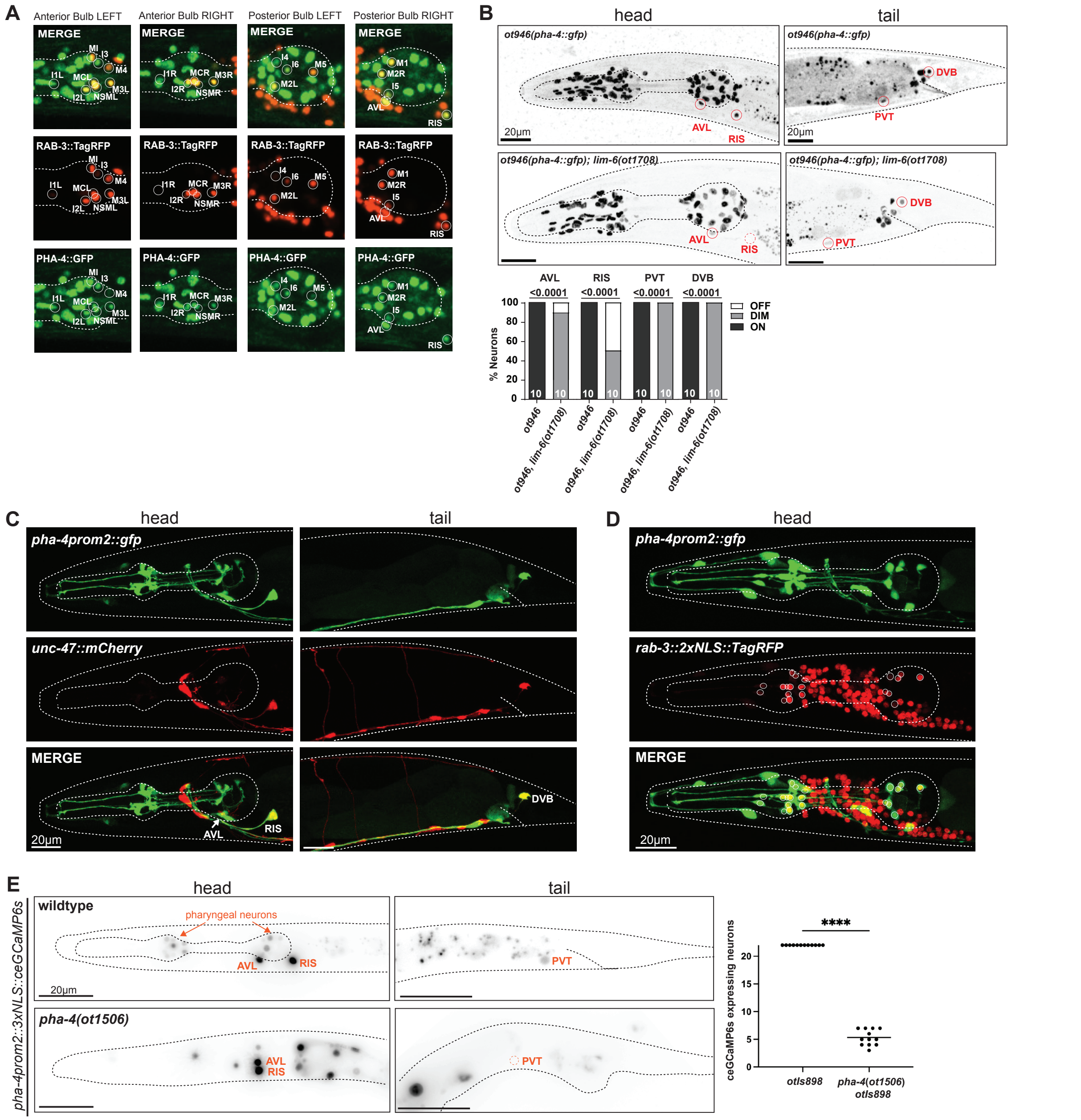


**Supplementary Figure 1: Expression of *pha-4* reporter reagents.**

**A:** *pha-4* expression (*ot946*) in PENs into adulthood. Representative images of a Day 1 Adult worm. *pha-4* expression (*ot946*) co-localizes with a neuron-specific reporter gene *rab-3* (allele) in all 20 neurons of the pharyngeal nervous system.

**B:** *lim-6(ot1708),* a molecular null allele with a complete locus deletion, results in a decrease in expression of the *pha-4(ot946)* reporter allele. Representative image of a Day 1 adult worm.

**C, D:** Overlap of *pha-4prom2::gfp* transgene (*otEx8085*) with the GABAergic marker *unc-47::mCherry* transgene (*otIs348,* panel C) and the panneuronal marker transgene *rab-3::2xNLS::TagRFP* (*otIs355,* panel D).

**E:** Expression pattern of *pha-4prom2* driving nuclear-localized GCaMP - *pha-4prom2::3xNLS::ceGCaMP6s::unc-54 3' UTR* (*otIs898*), in wildtype and *pha-4*(*ot1506)* null mutant animals. Data points representing each individual worm assayed are plotted, and the horizontal line in the middle of data points represents the median value of biological replicates. **** represents P < 0.0001 in Dunn's multiple comparison test after a Mann-Whitney test.

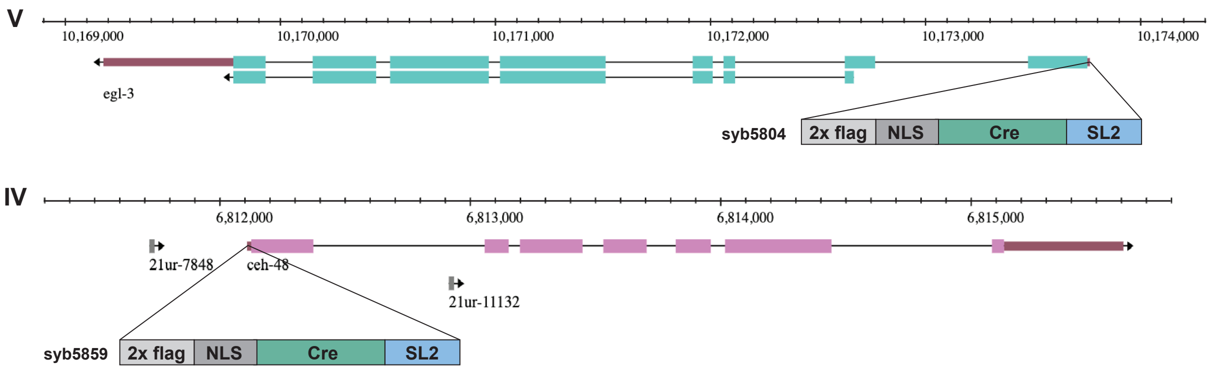


**Supplementary Figure 2: Locus schematic of *egl-3::Cre* and *ceh-48::Cre* drivers.**
**A:** *egl-3* locus schematic showing the *egl-3* CRISPR/Cas-9-engineered Cre insertion (*syb5804)*
**B:** *ceh-48* locus schematic showing the *ceh-48* CRISPR/Cas-9-engineered Cre insertion (*syb5859*).

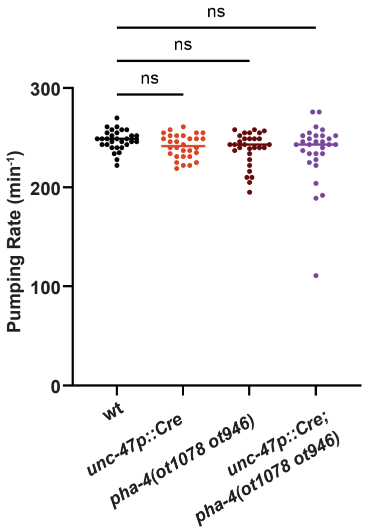


**Supplementary Figure 3: Characterization of effects of cell-specific *pha-4* removal.**
Effect of *pha-4* removal from AVL, RIS, and DVB on pumping behavior via cell-specific removal of *pha-4* using the floxed *pha-4* locus (*ot946 ot1078*) and *unc-47::Cre*(*arTi479*).

**Supp Table S1: Strain list.** A list of all strains used in this study.

| Strain number | Genotype |
|---|---|
| N2 | C. elegans Strain N2 (WormBase: WBStrain00000001) |
| GS10011 | arTi479 [unc-47p::cre::unc-54 3'UTR in miniMOS vector] I. |
| LSC27 | pdf-1(tm1996) III |
| NQ1208 | qnEx643 [Pflp-11::HisCl::SL2::mCherry, Pmyo-2::mCherry] |
| NY2080 | ynIs40 [flp-11p::GFP] V. |
| OH12496 | otIs518 [eat-4fosmid::SL2::cherry; pha-1(+)]; him-5(e1490) V |
| OH15039 | otIs625 [cat-1fos::SL2::mCherry::H2B; pha-1(+)] II; him-5(e1490) V |
| OH15568 | unc-17(ot907[unc-17::mKate2::3xFLAG] IV; him-5(e1490) V |
| OP171 | unc-119(ed3) III.; wgIs171 [egl-38::TY1::EGFP::3xFLAG(92C12) + unc-119(+)] |
| OP387 | unc-119(tm4063) III; wgIs387 [lim-6::TY1::EGFP::3xFLAG + unc-119(+)]. |
| PHX3208 | nlp-40(syb3208 [nlp-40::T2A::3XNLS::GFP]) I |
| PHX3330 | pdf-1(syb3330 [pdf-1::T2A::3XNLS::GFP]) III |
| PHX4861 | gpla-1(syb4861[flr-2::SL2::GFP::H2B]) V. |
| OH15876 | pha-4(ot946[pha-4::GFP::TEV::LoxP::3xFLAG]) V |
| OH16611 | otIs355 [rab-3::TagRFP] IV; pha-4(ot946[pha-4::GFP::TEV::LoxP::3xFLAG]) V |
| OH16691 | otIs355 [rab-3::TagRFP] IV; pha-4(ot946 ot1078[pha-4::GFP::TEV::LoxP::3xFLAG floxed]) V |
| PHX5755 | otIs355 [rab-3::TagRFP] IV; pha-4(ot946 ot1078 syb5755[pha-4::GFP::TEV::LoxP::3xFLAG::AID floxed]) V |
| OH17579 | oxIs12 [unc-47p::GFP] X. |
| OH18512 | otEx8085 [pha-4prom2::GFP::unc-54 3'UTR, pha-1(+)]; pha-1(e2123)III |
| OH18562 | unc-119(ed3) III; otIs898[pha-4prom2::3xNLS::ceGCaMP6s::unc-54 3' UTR] *oxTi553[eft-3p::tdTomato::H2B::unc-54 3'UTR + Cbr-unc-119(+ |
| OH18595 | unc-25(ot1372[unc-25::t2a:gfp::h2b]) III |
| OH18707 | otIs904[ges-1p::ins-1(genomic)::tagRFP::SL2::gfp::his-44::tbb-2 3’ UTR, inx-6p18::tagRFP::unc-54 3' UTR] *oxTi553[eft-3p::tdTomato::H2B:: |
| OH19200 | pha-4(ot1506) V. |
| OH19864 | fog-2(q71) pha-4(ot1506)/stu-3(q265) rol-9(sc148)V. |
| OH19201 | pha-4(ot1505 ot946[pha-4::gfp]) V. |
| OH19868 | fog-2(q71) pha-4(ot1505)/stu-3(q265) rol-9(sc148)V. |
| OH19280 | aex-5(ot1532[aex-5::SL2::gfp::his-44]) I. |
| OH19299 | unc-75(ot1539 [mScarlet-I3::unc-75]) I |
| PHX5804 | egl-3(syb5804[flag-NLS-Cre::SL2::egl-3]) V |
| PHX5859 | ceh-48(syb5859[flag-NLS-Cre::SL2::ceh-48]) IV |
| OH19848 | otIs946 [pha-4prom2::mNeonGreen::PH::p10 3'UTR, pha-4prom2::3xNLS::mScarlet-I3::unc-54 3'UTR] *oxTi553[eft-3p::tdTomato::H2B::unc- |
| OH19864 | otIs355 [rab-3::TagRFP] IV; fog-2(q71) pha-4(ot1506)/stu-3(q265) rol-9(sc148)V |
| OH19865 | arTi479 [unc-47p::cre-opti::unc-54 3'UTR in miniMOS vector] I; pha-4(ot946 ot1078 syb5755[pha-4::GFP::TEV::LoxP::3xFLAG::AID floxed]) |
| OH19866 | arTi479 [unc-47p::cre::unc-54 3'UTR in miniMOS vector] I; gpla-1(syb4861[flr-2::SL2::GFP::H2B]); pha-4(ot946 ot1078 syb5755[pha-4::GFP: |
| OH19868 | otIs355 [rab-3::TagRFP] IV; fog-2(q71) pha-4(ot1505 ot946[pha-4(intron1del)::GFP::TEV::LoxP::3xFLAG])/stu-3(q265) rol-9(sc148)V |
| OH19869 | otIs696 [NeuroPAL]; pha-4(ot946 ot1078 syb5755[pha-4::GFP::TEV::LoxP::3xFLAG::AID floxed] V |
| OH19870 | ynIs40 [flp-11p::GFP]; pha-4(ot1506) V. |
| OH19871 | gpla-1(syb4861[flr-2::SL2::GFP::H2B]); pha-4(ot1506) V. |
| OH19873 | aex-5(ot1532[aex-5::SL2::gfp::his-44]) I; pha-4(ot1506) V. |
| OH19875 | nlp-40(syb3208 [nlp-40::T2A::3XNLS::GFP]) I; pha-4(ot1506) V. |
| OH19879 | ceh-48(syb5859[flag::NLS::Cre::SL2::ceh-48]) IV; pha-4(ot946 ot1078 [pha-4::GFP::TEV::LoxP::3xFLAG floxed]) V |
| OH19888 | otIs904[ges-1p::ins-1(genomic)::tagRFP::SL2::gfp::his-44::tbb-2 3’ UTR, inx-6p18::tagRFP::unc-54 3' UTR] *oxTi553[eft-3p::tdTomato::H2B:: |
| OH19943 | unc-75(ot1539 [mScarlet-I3::unc-75]) I; egl-3(syb5804[flag::NLS::Cre::SL2::egl-3]); pha-4(ot946 ot1078 [pha-4::GFP::TEV::LoxP::3xFLAG flo |
| OH19945 | pha-4(ot946 ot1078 syb5755[pha-4::GFP::TEV::LoxP::3xFLAG::AID floxed]; otIs908[pha-4prom2::TIR1(F79G)::mTurq2::tbb-2 3’ UTR, unc-1 |
| OH19946 | arTi479 [unc-47p::cre::unc-54 3'UTR in miniMOS vector] I.; pha-4(ot946 ot1078 syb5755[pha-4::GFP::TEV::LoxP::3xFLAG::AID floxed]) V.; [illegible] |
| OH20089 | otEx8333 [sre-22p::gfp::h2b::tbb-2 3'UTR, unc-122p::gfp] |
| OH20090 | otEx8334 [sre-22p::gfp::h2b::tbb-2 3'UTR, unc-122p::gfp] |
| OH20091 | otEx8335 [sre-22p::gfp::h2b::tbb-2 3'UTR, unc-122p::gfp] |
| OH20092 | otEx8336 [sre-22p::HisCl::SL2::gfp::tbb-2 3'UTR, unc-122p::gfp] |
| OH20093 | otEx8337 [sre-22p::HisCl::SL2::gfp::tbb-2 3'UTR, unc-122p::gfp] |
| OH20095 | otEx8339 [sre-22p::2xFlag::NLS::Cre::unc54 3'UTR, unc-122::gfp]; egl-3(nu1711[egl-3 floxed]) V |
| OH20096 | otEx8340 [sre-22p::2xFlag::NLS::Cre::unc54 3'UTR, unc-122::gfp]; egl-3(nu1711[egl-3 floxed]) V |
| OH20097 | otEx8341 [sre-22p::2xFlag::NLS::Cre::unc54 3'UTR, unc-122::gfp]; egl-3(nu1711[egl-3 floxed]) V |
| OH20098 | otEx8342 [sre-22p::HisCl::SL2::gfp::tbb-2 3'UTR, unc-122p::gfp] |
| OH20099 | otEx8343 [sre-22p::2xFlag::NLS::Cre::unc54 3'UTR, unc-122::gfp]; pha-4(ot946 ot1078[pha-4::gfp-loxP]) V |
| OH20100 | otEx8344 [sre-22p::2xFlag::NLS::Cre::unc54 3'UTR, unc-122::gfp]; pha-4(ot946 ot1078[pha-4::gfp-loxP]) V |
| OH20101 | otEx8345 [sre-22p::2xFlag::NLS::Cre::unc54 3'UTR, unc-122::gfp]; pha-4(ot946 ot1078[pha-4::GFP::TEV::LoxP::3xFLAG floxed]) V |
| OH20103 | otIs518 [eat-4fosmid::SL2::cherry; pha-1(+)] II; egl-3(syb5804[flag::NLS::Cre::SL2::egl-3]); pha-4(ot946 ot1078 [pha-4::GFP::TEV::LoxP::3x |
| OH20104 | otIs625 [cat-1fos::SL2::mCherry::H2B; pha-1(+)] II.; egl-3(syb5804[flag::NLS::Cre::SL2::egl-3]); pha-4(ot946 ot1078 [pha-4::GFP::TEV::Lox |
| OH20105 | otIs518 [eat-4fosmid::SL2::cherry; pha-1(+)] II.; pha-4(ot1505 ot946[pha-4::gfp]) V. |
| OH20106 | otIs625 [cat-1fos::SL2::mCherry::H2B; pha-1(+)] II.; pha-4(ot1505 ot946[pha-4::gfp]) V. |
| OH20107 | unc-17(ot907[unc-17::mKate2::3xFLAG] IV; pha-4(ot1505 ot946[pha-4::gfp]) V. |
| OH20108 | wgIs387 [lim-6::TY1::EGFP::3xFLAG + unc-119(+)].; pha-4(ot1506) V. |
| OH20109 | wgIs387 [lim-6::TY1::EGFP::3xFLAG + unc-119(+)].; pha-4(ot1505 ot946[pha-4::gfp]) V. |
| OH20110 | arTi479 [unc-47p::Cre::unc-54 3'UTR in miniMOS vector] I; unc-25(ot1372[unc-25::t2a:gfp::h2b]) III; pha-4(ot946 ot1078 syb5755 [pha-4::G |
| OH20111 | nlp-40(syb3208 [nlp-40::T2A::3XNLS::GFP]) I; pha-4(ot1505 ot946[pha-4::gfp]) V. |
| OH20113 | wgIs171 [egl-38::TY1::EGFP::3xFLAG(92C12) + unc-119(+)]; pha-4(ot1506) V |
| OH20114 | pdf-1(syb3330 [pdf-1::T2A::3XNLS::GFP]) III.; pha-4(ot1506) V. |
| OH20115 | otEx8333 [sre-22p::gfp::h2b::tbb-2 3'UTR, unc-122p::gfp]; pha-4(ot1506) V. |
| OH20116 | otIs518 [eat-4fosmid::sl2::mcherry; pha-1(+)] II.; pha-4(ot946 ot1078 syb5755[pha-4::GFP::TEV::LoxP::3xFLAG::AID floxed]; otIs908[pha-4 |
| OH20241 | otEx8356 [pha-4prom1::GFP::unc-54 3'UTR, pha-1(+)]; pha-1(e2123)III |
| OH20243 | otIs348[unc-47p(300bp)::mCherry::unc54-3'UTR, pha-1(+)] IV; pha-4(ot946[pha-4::gfp]) |
| OH20244 | otIs355 [rab-3::TagRFP] IV; otEx8085 [pha-4prom2::GFP::unc-54 2'UTR, pha-1(+)] |
| OH20245 | otIs348[unc-47p(300bp)::mCherry::unc54-3'UTR, pha-1(+)] IV; fog-2(q71) pha-4(q490)/stu-3(q265) rol-9(sc148)V |
| OH10598 | otIs348[unc-47p(300bp)::mCherry::unc54-3'UTR, pha-1(+)] IV. |
| ZW1273 | homt-1(zw94) I |
